# Circadian immunometabolic states impart a temporal response to SARS-CoV-2 spike proteins in mammalian macrophages

**DOI:** 10.64898/2026.02.24.707668

**Authors:** Sharleen M. Buel, Jayashree Balaraman, Meaghan S. Jankowski, Kim Hixson, Yuqian Gao, Young-Mo Kim, Nathalie Muñoz, Jennifer E. Kyle, Mary S. Lipton, Carrie D. Nicora, Paul D. Piehowski, Scott E. Baker, Jennifer M. Hurley

## Abstract

Circadian rhythms, driven by 24-hour molecular oscillators, or “clocks”, widely tune physiology to the daily rhythms of light and dark to enhance organismal fitness. In mammals, the cellular immune response is tightly regulated by these rhythms such that immunometabolic output is coordinated across the day, consolidating macrophage physiology into temporally distinct phases that determine the macrophage response to stimuli. Importantly, key proteins in the macrophage response to viral infection have been found to be under circadian control, and time of day of adjuvant application is known to affect the efficacy of vaccinations, including in the case of the SARS-CoV-2 virus. However, little is known about the molecular changes that underly the temporal response to vaccine application. Therefore, to investigate the circadian response of macrophage physiology to adjuvant exposure, we exposed primary mouse and human macrophages to the SARS-CoV-1 and CoV-2 spike proteins at different times over the circadian day. To further explore the time-of-day effect, we performed a multi-omics analysis and *in vitro* tissue culture assays examining macrophage responses over circadian time. We found that, conserved across the species, the timing of spike protein exposure dictated two distinct temporal responses which were characterized by hallmarks of immunometabolic suppression and modest immunometabolic activation. Intriguingly, these temporal responses were driven by central metabolic and mitochondrial changes rather than classical immune activation, suggesting immunometabolic control is a primary regulator of the temporal response of immune cells to stimuli.

## Introduction

Macrophages are a dynamic cell type that play important protective roles within the innate immune system, comprising approximately 10% of all lymphocytes within the human body [1]. Macrophages function as the first line of immune defense against pathogens, orchestrating inflammatory reactions and activating the adaptive immune system through multiple signaling pathways [2]. In the response to COVID-19 Severe Respiratory Syndrome (SARS-CoV-2), macrophages are essential in mediating hyperinflammation and cytokine levels to contribute to disease progression and vaccine response [3,4]. The importance of the inflammatory response in macrophages in the context of SARS-CoV-2 underscores the need to understand the cellular mechanisms underlying macrophage inflammatory activation and the processes that regulate this activation.

Central metabolism is a key driver of inflammatory activation as well as homeostatic maintenance in macrophages. Whether activated or naïve, macrophages must use energy to migrate throughout resident tissues and sample the cellular milieu (via phagocytosis and internal processing) [5]. These cellular functions are metabolically expensive and require high energy levels, mainly generated by the central metabolic pathways of glycolysis, the TCA cycle, and oxidative phosphorylation (OXPHOS) [2]. To mount a pro-inflammatory response, naïve macrophages that encounter pathogens must extensively rewire central metabolic pathways to quickly increase the production of energy. This rewiring enhances glycolysis to expand the energy output to support pathogen intake, processing, and greater immune activation [6]. In concordance with this, macrophages downregulate enzymes within the TCA cycle, allowing for accumulation of certain TCA metabolites, halting the flow of metabolites and electrons into the OXPHOS chain, and exchanging the production of ATP for that of reactive oxygen species (ROS) to support elements of pro-inflammatory activation [6,7]. To further support this metabolic reprogramming, mitochondria, which house these metabolic reactions, undergo fragmentation, effectively decreasing ATP generation by the ATP synthase complex [7]. Mitochondrial fragmentation also acts as an immune signaling platform, where ROS, bioactive lipids, and changes in MMP are transduced to enhance biological signals [8].

Given the importance of central metabolism and mitochondrial dynamics in both homeostatic functioning and the pro-inflammatory response in macrophages, these processes are tightly regulated. Circadian rhythms, the rhythms that tune physiology to the 24hr day/night cycle, have been shown to participate in this regulation. The core circadian clock that tunes this 24-hr cycle is comprised of an activating complex, BMAL1/ARNTL and CLOCK, and a repressive complex, PER and CRY, that time cellular functions across the central dogma. This resultant circadian regulation has been widely shown to time metabolic pathways, including central metabolic enzymes, immune factors, and mitochondrial dynamics across many cell types [9]. Specifically, molecular analysis shows that as much as ∼30% of the proteome in peripheral macrophages is tuned to oscillate with a circadian rhythm [10]. This oscillation supports temporal immune function and appears to organize macrophage function into distinct pro- and anti-inflammatory phases, driven by changes to the immunometabolic state [10,11]. However, despite the evidence of significant circadian regulation on important immunometabolic functions, little is known about the resultant effect on the metabolites themselves and how the availability of these central metabolic components affects the macrophage response to stimuli.

This immunometabolic response is important, as many pathogens target and disrupt central metabolism and mitochondrial components during infection [12]. For example, exposure to SARS-CoV-2 causes widespread disruption of central metabolic pathway proteins and metabolites in addition to mitochondrial function, which all ultimately assist in viral replication [13]. However, little has been done to examine central metabolic responses in macrophages following exposure to SARS-CoV-2. Moreover, while the effectiveness of COVID vaccination is known to vary with the time of day of administration, there has been no work examining the connection between circadian immunometabolic regulation and the response to SARS-CoV-2 [14]. What is known is that the macrophage response is in part due to the interaction of macrophage cell surface receptors with the SARS-CoV-2 spike protein, which decorates the outer surface of the virus, facilitating cellular entry [15]. In fact, exposure to just the S1 subunit of the spike protein (amino acid residues 14-685) is sufficient to disrupt mitochondrial morphology and glycolysis in macrophages and other cell types [16–19]. This metabolic reprogramming persists after exposure, causing hyperinflammation and dysregulated immune responses associated with long COVID [17].

Given the above, we hypothesized that exposure to the SARS-CoV spike protein elicits a differential macrophage response based on the time of day of exposure. To investigate how the macrophages’ pathogen responses were altered depending on the time of day they encountered SARS-CoV spike protein, we worked with SARS-CoV-1 and CoV-2 spike proteins in both wild-type mouse and human primary macrophages. We found consistent circadian timing of immunometabolic regulation in both species that resulted in similar temporal response signatures to spike protein exposure. These responses were primarily driven by changes to central metabolic pathways and mitochondria—with little contribution by classical inflammatory pathways—that were different depending on the timing of spike protein treatment. Further, we found there were widespread time-of-day specific physiological responses, including temporal changes to protein, metabolite, and lipid levels. In total, our data suggest the clock coordinates two distinct temporal immune responses to COVID spike proteins in macrophages, an immunometabolic repression phase and a modest immunometabolic enhancement phase, demonstrating a physiologically relevant role for circadian immunometabolic timing in the response to SARS-CoV-2.

## Results

### Analysis of the murine macrophage circadian proteome, metabolome, and lipidome reveals circadian control across multiple molecular levels

To understand the effects of time-of-day of spike protein exposure on the macrophage molecular response, we first had to understand the basal circadian changes elicited by the clock on macrophage physiology. We have previously explored the circadian control of proteomics in mouse BMDMs, demonstrating extensive timing of metabolism and the immune response at the cellular level [10,20]. However, an analysis of how circadian regulation affects the individual metabolites that participate in these cellular functions has yet to be completed. Therefore, we first aimed to identify circadian regulation of metabolic elements in mammalian macrophages. To do so, we isolated, differentiated, and circadianly synchronized mouse bone marrow-derived macrophages from male PER2::LUC mice as previously described **(Fig 1A),** using luminescence traces to confirm our protocol resulted in ∼24-h rhythms as we have done previously [10]. To avoid the known artifactual gene expression that occurs immediately following serum shock, we began sampling the time course at 22 hours post-serum shock (HPS22), collecting and freezing cell pellets every 4 hours for 24 hrs [21]. The MPLEx method was then used to extract total proteins, metabolites, and lipids from each sample [22] **(Fig 1A)**. By using the MPLEx approach, the same BMDM cell pellet could be analyzed for total proteomics, metabolomics, and lipidomics, thereby creating a matching data set for each timepoint, which could be used to perform coordinated downstream multi-omics analyses.

**Fig 1.**
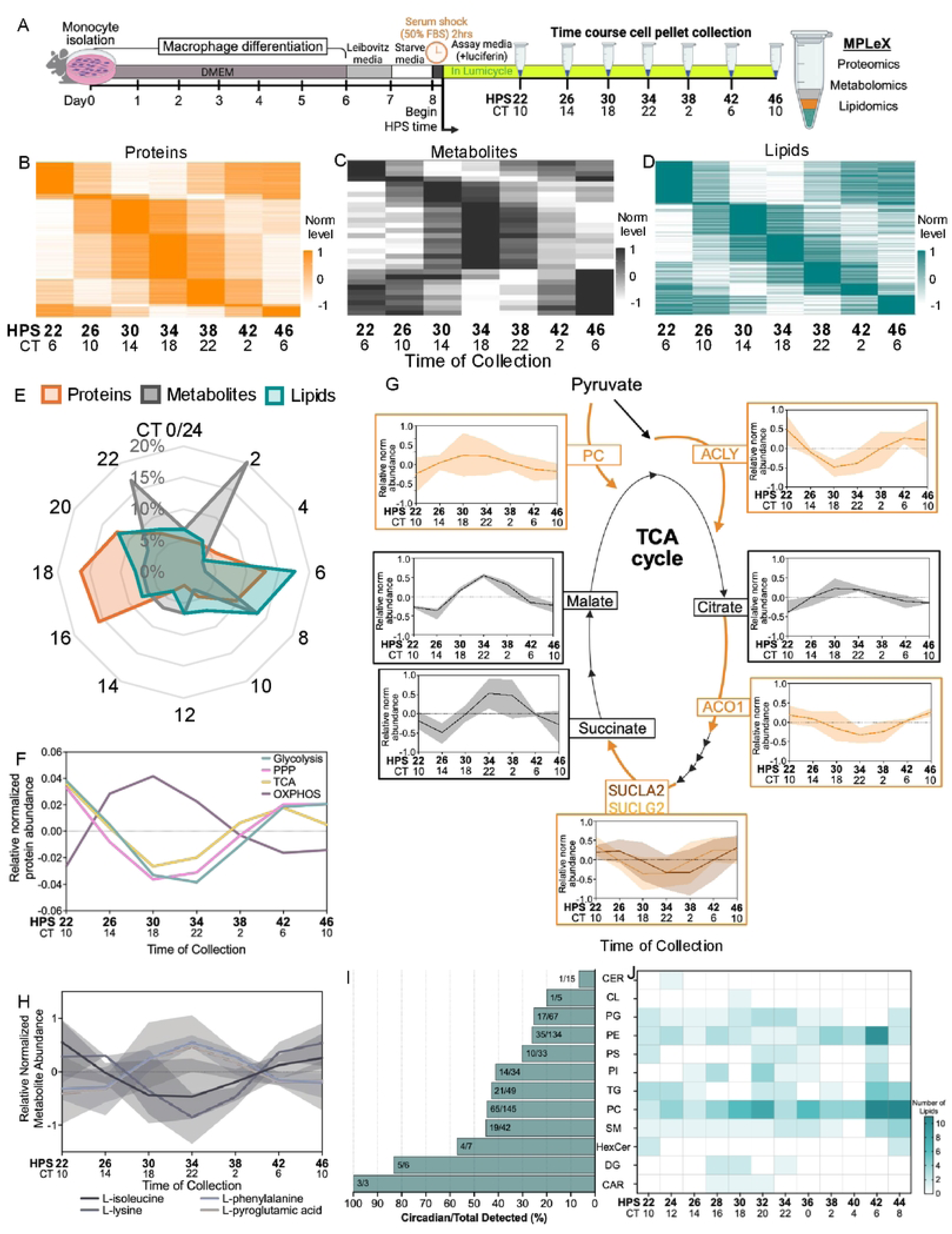
Circadian rhythms control the proteins, metabolites, and lipids of central metabolism and the inflammatory response. **(A)** Schematic of the timeline for collecting multi-omics data from circadianly-synchronized mouse bone marrow-derived macrophages (mBMDMs). **(B-D)** Heatmaps representing relative high to low abundances of circadian **(B)** proteins (high: orange, low: white), **(C)** metabolites (high: gray, low: white), and **(D)** lipids (high: teal, low: white) with significant circadian oscillations (Period: 20-28 hours, BH-adj p-val <0.05) as identified by ECHO [23]. **(E)** The percentage of total oscillating proteins (orange), metabolites (grey), and lipids (teal) peaking at a given time plotted on a radial histogram by circadian time of peak phase, binned in 2-h windows. **(F)** ECHO fitted models to the normalized relative abundances of all proteins within the listed central metabolic pathways. **(G)** Circadianly oscillating proteins (orange) and metabolites (gray) that are involved in the TCA cycle plotted in the context of the pathway. ECHO fitted model shown in dark line and ±1SD of the data shown in shaded portion. **(H)** ECHO fitted models of normalized relative abundances for circadianly oscillating amino acids that are active in inflammation and oxidative stress [10]. Shaded regions indicate ±1SD of the data. **(I)** Proportion of circadian lipids over total detected lipids in each lipid class (numbers of circadian/total). **(J)** A heat map comparing the peak phases of oscillating lipids by lipid class [10]. Ceramides (Cer), cardiolipins (CL), phosphatidylglycerols (PG), phosphatidylethanolamines (PE), phosphatidylserines (PS), phosphatidylinositols (PI), triglycerides (TG), phosphatidylcholines (PC), sphingomyelins (SM), hexocylceramides (HexCer), diacylglycerols (DG), acylcarnitines (CAR), HPS = hours post-synchronization, CT = circadian time.

To perform the total proteomics analysis, randomized 16-plex TMT-labeled MS/MS analyses were run on samples taken from the interphase MPLEx layer. In total, 180,261 unique peptides across 14,025 proteins were detected in the native macrophage conditions. We then used LIMBR to impute missing data (if <30%) and remove batch effects. Finally, we rolled the peptides up to the protein level by averaging their relative abundances, identifying 7,331 total proteins that were reliably detected in our data set [24]. Correlation plots and PCA analyses of these data indicated the biological replicates were highly correlated, with most variance captured in principal components 1 and 2 **(S1A-F Fig)**. We next modeled the rhythms of the proteome using the tool ECHO [23]. We first looked at the core clock proteins that were detected in the dataset, including BMAL1, PER1, and CRY1, and auxiliary loop repressor NR1D2 (Rev-Erbβ). Due to the canonically low levels of these proteins, we analyzed their oscillations pre-LIMBR [10]. We found these proteins oscillated with periods ranging from 20-28 hours with similar patterns as were seen previously **(S1G Fig)** [10].

To gather data on total metabolomics, we subjected metabolites from the upper aqueous phase of the MPLEx layers to GC-MS/MS. An internal standard (13C-labeled fumaric acid) was additionally run alongside our samples for further validation. Our methods detected 230 metabolites which were then subjected to processing by LIMBR, a circadian-specific processing program which can impute missing data and remove batch effects without conflating circadian trends with noise. Processing our metabolomics data using LIMBR returned 226 metabolites that were reliably detected (i.e. >70% coverage), which were used for circadian analysis. To quantify lipids within our samples, we subjected the bottom organic phase of the MPLEx layers to LC-ESI-MS/MS in both positive and negative modes. 540 total lipids (286 in negative mode, 254 in positive mode) with 100% coverage were detected between both positive and negative runs and separately sent through LIMBR processing before being used for circadian analysis.

We next analyzed each of our data sets using ECHO to determine the range of circadian timing within the proteome, metabolome, and lipidome in macrophages [23]. Based on the range of periods of the core clock proteins and the 4-hour resolution of the omics data, a protein, metabolite, or lipid was defined as circadianly oscillating if it had a period within the range of 20-28 hours and a BH-adj p-value of <0.05. These parameters identified 2,622 (35.8%) proteins, 30 (13.3%) metabolites, and 193 (35.7%) lipids that oscillated with a circadian period **(Fig 1B-D)**.

### Metabolic regulation is highly circadianly coordinated across macrophage molecular features

To understand how the circadian clock orchestrates the overall architecture of rhythms across these three omics data sets over the circadian day, we graphed the circadian timing (CT) of their peaks. We calculated CT by adjusting all circadian protein, metabolite, and lipid periods to 24 hours and aligned our *in vitro* PER2 oscillations in our proteomics data to previous *in vivo* PER2 oscillations [10,11]. Therefore, we were able to determine that HPS16 = CT4. Overlaying the peak circadian timings of each omics data type on a 24-hour clock-style graph revealed distinct patterns of peak timing for each data type. Proteins generally followed the bimodal pattern we identified previously, in this case with a slight shift in peak during the anti-inflammatory and inflammatory phases (∼CT6 and ∼CT18) as compared to [10]. The peak in oscillations of the lipids also segregated into two peaks (∼CT6 and ∼CT20) while the metabolites had three distinct peaks in oscillations (CT2, CT8, CT22) **(Fig 1E)**.

We next subjected the proteomics data to Gene Ontological analysis via KEGG pathways, to align the data with our previously published data set, revealing consistencies, namely in the control of central metabolic pathways [10,25]. Glycolysis, the pentose phosphate pathway (PPP), the TCA cycle, and oxidative phosphorylation (OXPHOS) were all shown to be highly circadianly regulated (14-20% of the detected pathway) (**Supplemental Dataset**). We next modeled the normalized protein abundances of all detected pathway proteins and identified that each pathway showed a significant circadian oscillation in overall levels **(Fig 1F, S1H-K Fig)**. To further understand circadian regulation of the OXPHOS pathway, we identified the circadianly-controlled proteins in the Electron Transport Chain (ETC) Complexes I, II, IV, and V (as delineated in the OXPHOS pathway in the KEGG database for *M. musculus—*mmu00190 [25]), demonstrating that these proteins oscillate in phase with each other, peaking around HPS30/CT18 **(S1L-O Fig)**. These data suggest the resultant central metabolites should also oscillate with a circadian period.

To determine how oscillations in central metabolic proteins affected metabolite levels, we next assigned the detected metabolites to their respective metabolic pathways. We found few detected metabolites within glycolysis or the pentose phosphate pathway. However, we did observe metabolites within the TCA cycle that play key roles in macrophage metabolic rearrangement under polarizing (pro- and anti-inflammatory) conditions in our data set [2]. We found many of these metabolites oscillated with a circadian period, including citric acid, succinate, and malate. When mapped onto the TCA cycle pathway, we noted the enzymes acting on these metabolites were also under circadian control **(Fig 1G)**, illustrating direct regulation by the circadian clock at key points of metabolic control. Beyond central metabolism, we found several other classes of metabolites that oscillated with a circadian period. One of these highly oscillatory classes was amino acids, including isoleucine, lysine, phenylalanine, and pyroglutamic acid, which are all associated with inflammation and oxidative stress **(Fig 1H)** [26–30].

Beyond the metabolites, we also investigated oscillations within the lipidome. When we broke down the circadian lipids by class, we found the highest levels of circadian regulation were seen in acylcarnitines (CAR), diacylglycerides (DG), hexosylceramide (HexCer), sphingomyelins (SM), phosphatidylethanolamines (PE) and phosphatidylcholines (PC) **(Fig 1I)**. We noted that the peak phase of both PC and PE classes, which ensure stability of OXPHOS complexes within the inner membrane, arose between CT2-10 **(Fig 1I)** [31]. The phase timing of additional mitochondrial-related lipid classes PC, CAR, and cardiolipins (CL) coincides with the rise of OXPHOS complex components **(Fig 1F and J)**, suggesting circadian regulation may additionally coordinate OXPHOS through the timing of these lipids [32,33].

### Macrophages have time of day immune-related responses to SARS spike proteins

Having demonstrated the multi-layered circadian regulation of immunometabolic pathways in macrophages, this data further suggested there may be time-of-day ramifications to the response to an exogenous stimulus. To test this theory, we again isolated and synchronized mBMDMs from young, male PER2::LUC mice, and then exposed them to the S1 subunit of three different Severe acute respiratory syndrome (SARS) coronaviruses at seven different time points across the circadian day (HPS16, 20, 24, 28, 32, 36) for 6 hrs **(Fig 2A)**. We chose the S1 spike proteins from SARS-CoV-1, WT SARS-CoV-2, and SARS-CoV-2^D614G^ **(S2A Fig)** as these regions contain the binding domain for the ACE2 receptor in murine macrophages and have been shown to be an important factor in the response to SARS infection [34]. The spike-protein exposed mBMDMs were then analyzed using multi-omics analysis as described above. After cleaning and pre-processing the data sets using LIMBR [24], we assessed the validity and quality of the data. Correlation coefficients demonstrated that the biological replicates of all spike protein treatments were highly correlated within each proteomic, metabolomic, and lipidomic data sets and clustered independently in the PCA space **(S1A-F Fig)**.

**Fig 2.**
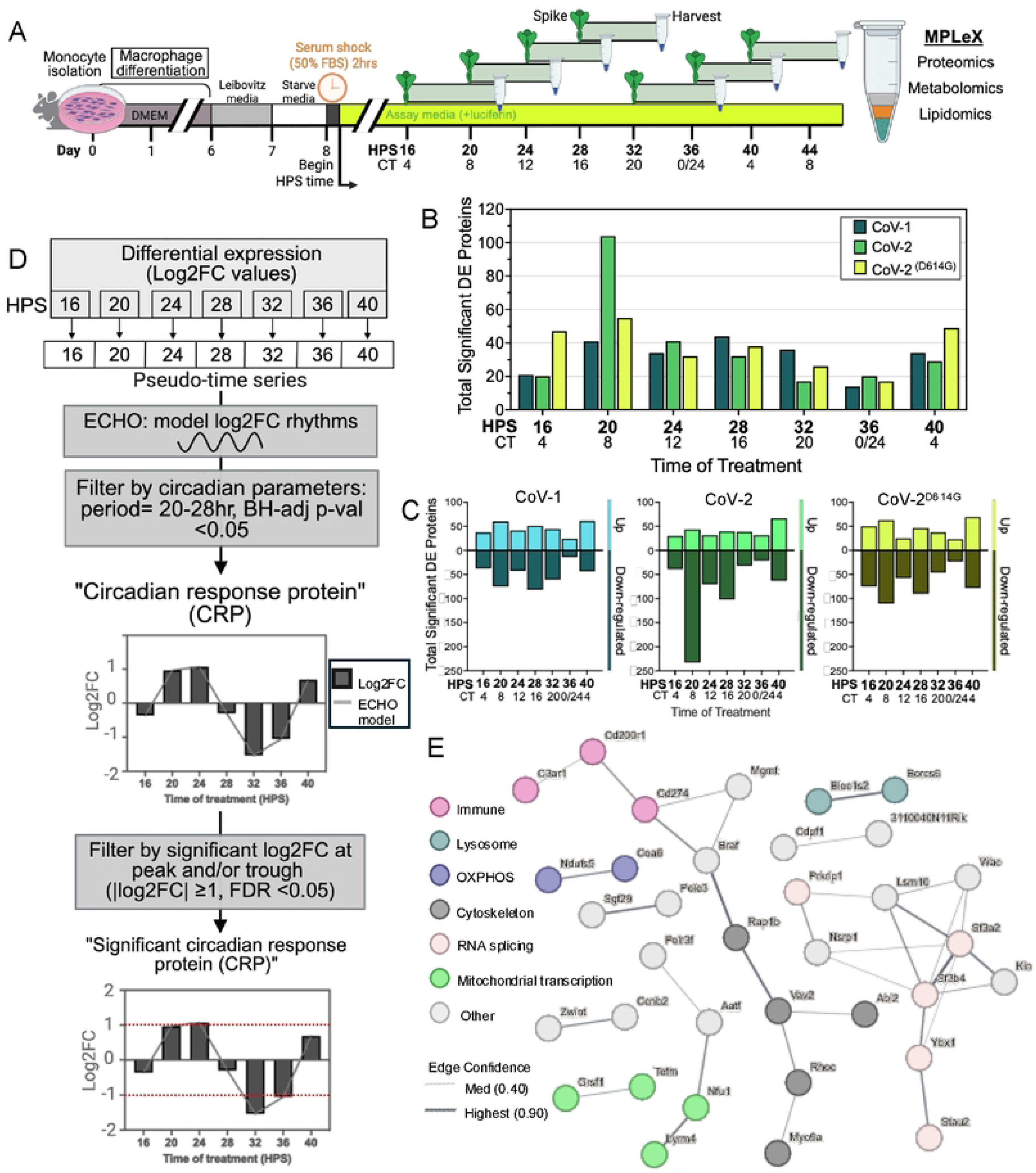
Circadian rhythms regulate inflammatory and immune responses to SARS spike proteins. **(A)** A schematic of the timeline of the experimental setup for circadian synchronization of mouse bone marrow-derived macrophages (mBMDMs), exposure to SARS spike proteins, and collection of samples for multi-omics processing. **(B)** A plot of the number of significant Differentially Expressed (DE) proteins at each time point, for each spike protein treatment. **(C)** The number of significant up- and down-regulated proteins at each time point, for each spike protein treatment. **(D)** A flow diagram describing how we defined circadian response proteins (CRPs) after spike protein treatment. First, a pseudo-timecourse was created for each protein by joining together log2FC values from each time point (e.g. HPS16, 20, 24, 28, 32, 36, 40). This pseudo-timecourse was then modeled using ECHO with circadian period and BH-adj p-val cutoffs [23]. To detect CRPs with a significant peak or trough of response, we determined peak and trough times from the ECHO model and identified the corresponding log2FC value at these time points. If the log2FC value at either or both time points met our log2FC cutoff values (dotted red lines shown on bottom example graph), we categorized it as a significant CRP. **(E)** The stringDB (v12.0) network of significant circadian response proteins (CRPs). Node color indicates functional category for a given protein, as delineated in the key. Edges, or lines, represent both functional and physical protein associations, and the edge width represents the strength of interaction evidence, as delineated in the key.

To understand how the time of day affected the molecular components of the cell, we next analyzed each omics set for differential levels relative to the vehicle-treated control using the edgeR package in R [35]. Due to limitations within edgeR and other differential expression packages for time-series data with treatments, we compared each set of spike-treated samples separately by time point. We filtered these data using an FDR <0.05 and |log2FC| values of ≥ 1.0 to capture the significantly differentially-expressed proteins (DEPs), metabolites (DEMs), and lipids (DELs). We began by analyzing the overall change in the protein response by time of day. We found that, regardless of spike variant, treatment at HPS36 (CT0/24) induced the fewest significantly DEPs **(Fig 2B)** and treatment at HPS20 (CT8) displayed the highest number of significantly DEPs, which was primarily driven by the downregulation of proteins at HPS20 (CT8) **(Fig 2C)**.

Given the time-of-day specific response to spike protein exposure, we hypothesized proteins might exhibit circadian rhythms in their log2FC response to spike proteins, meaning they showed a differential expression to control that had a circadian rhythm. To identify proteins with a circadian response to spike protein exposure, we next stitched together each protein’s log2FC values from all treatment time points across the entire time series, creating a pseudo time-series that we modeled using ECHO [23] **(Fig 2D)**. We then filtered the ECHO output for the same circadian parameters as above (e.g. Period=20-28, BH-adj p-val <0.05) to identify “circadian response proteins” (CRPs). Although we identified thousands of CRPs, many of these CRPs had low log2FC values, suggesting a low biological effect of these CRPs. Therefore, we further filtered the CRPs for proteins with a peak or trough value of |log2FC| ≥ 1 and an FDR of <0.05 **(Fig 2D)**. This filtering identified 42, 47, and 15 CRPs in CoV-1, CoV-2, and CoV-2^D614G^ treatments, respectively. A StringDB network analysis of these CRPs revealed the interconnected terms were related to mitochondrial transcription, lysosome components, OXPHOS complexes, immune-related proteins, cytoskeleton rearrangement, and spliceosome components **(Fig 2E)** [36]. Taken together, this indicates that the proteins which respond circadianly to spike protein are important for some immune functions and broad regulatory pathways and shows clear evidence that there is an overall time-of-day specific response to the SARS spike proteins in macrophages.

### The core circadian clock is not affected by SARS spike protein exposure

Classical macrophage polarization has been shown to alter the period and amplitude of the core circadian clock [37,38]. A change to the core clock could explain the time-of-day specific protein responses we noted above. Therefore, as part of the above-described time course, we also tracked circadianly-synchronized mBMDMs exposed to the three SARS spike proteins at the same seven time points using a Lumicycle, quantifying luciferase—and thereby PER2::LUC expression—in response to spike protein exposure over 96 hrs. Surprisingly, using ECHO analysis we noted no changes to PER2 period, amplitude, or phase **(S2B, C Fig)** as compared to treatment by the classical stimuli LPS and IL-4, which produce marked phase shifts **(S2D Fig, Supplemental Dataset)**. We next looked at the other components of the core clock by tracking differential protein levels, from the proteomic analysis, of the detected clock proteins BMAL1/ARNTL, PER1, NR1D2 and casein kinase 1 isoform delta (CK1δ). Although we were unable to track changes in oscillation patterns, we found PER1 and CK1δ levels were significantly affected by exposure to CoV-1 and CoV-2 spike proteins at certain times (PER1 was upregulated at HPS28/CT16 by CoV-1 spike, while CK1δ was downregulated by both CoV-1 and CoV-2 at multiple time points (HPS20/8, HPS28/CT14, HPS32/CT20) **(S2E Fig)**. As both PER1 and CK1δ play roles in immune and metabolic regulation, this could be a potential avenue for differential output regulation [39,40]. However, as these changes were small, these data demonstrated that spike-protein specific CRPs likely do not stem from changes to the oscillation of the core circadian clock proteins.

### Macrophage cytokine release demonstrated minimal rhythmic responses to spike proteins

As the proteomic data suggested time-dependent immune activation to spike proteins, and the inflammatory response is an important driver of COVID pathology [41], we next evaluated the effect of the spike proteins on downstream immune output by tracking cytokine secretion. To assay the release of cytokines at different treatment times, we derived macrophages from PER2::LUC male mice and synchronized their circadian rhythms as described above. At the same 7 treatment times, we exposed our macrophages to either the vehicle control (PBS) or one of the SARS spike proteins (CoV-1, CoV-2, or CoV-2^D614G^). After 6 hours of incubation, we collected the supernatants and then used an ELISA to quantify concentrations of the pro-inflammatory cytokines (IL-6 and TNF-α), and anti-inflammatory cytokines (IL-10 and TGFβ-1). Under non-stimulatory conditions, IL-6, IL-10, and TNFα were constitutively released (e.g. not circadian) **(S3 A-D Fig)**. Treatment with some of the spike proteins did moderately increase the secretion of the pro-inflammatory cytokines IL-6 and TNFα, but these increases did not have a temporal trend. One anti-inflammatory cytokine, TGFβ-1, was released in a circadian manner in response to CoV-1 spike protein, with the peak around treatment time HPS32/CT20 (the end of the pro-inflammatory phase) **(S3E Fig)**. However, although we noted a cytokine response to many spike proteins, the overall measured levels were markedly lower than treatment with LPS and IL-4, suggesting moderate inflammatory activation output **(S3F, G Fig)**. These data demonstrated a limited rhythmic cytokine response to spike proteins, suggesting that time of day of exposure to the spike proteins does not have a significant circadian effect on cytokine levels.

### The circadian response to spike protein exposure aligns with central metabolite and metabolic pathway oscillations

As there was a limited temporal effect of spike-protein exposure on the classical immune output pathways in macrophages, we next tracked other facets of the macrophage response to determine other pathways affected by these CRPs that could play a role in the temporal response of macrophages to stimuli. Beyond the regulation of the release of cytokines, there are other ways to regulate the immune response of a macrophage. One key element of control lies in the regulation of central metabolism, which impacts macrophage physiology to tune anti-inflammatory/pro-inflammatory polarization, a process termed immunometabolism [42]. Immunometabolism is circadianly controlled, and in support of this we found several central metabolic pathways—namely glycolysis, TCA cycle, and oxidative phosphorylation (OXPHOS)—were under circadian control in our dataset **(Fig 1F-G, S1H-O)**.

Therefore, we next analyzed the effect of SARS spike proteins on central metabolic pathways based on the time of day of exposure. To do so, we modeled the log2FC values of all proteins detected in glycolysis, the TCA cycle, and oxidative phosphorylation (OXPHOS) by pathway using ECHO [23]. We found that in all spike protein treatments, the aggregate of the proteins in glycolysis responded in a circadian manner, with a peak around treatment time HPS28/CT16 (**Fig 3A, S3H Fig**). For the proteins in the pentose phosphate pathway (PPP), treatment with CoV-1 and CoV-2D^614G^ led to a circadian response, while for the proteins in the TCA cycle, only treatment with CoV-2 led to a circadian response (**Fig 3A; SI-J).** Contrary to the other metabolic pathways, the proteins in the OXPHOS pathway did not respond in a circadian manner to any treatment (**Fig 3A-C, S3K Fig**).

**Fig 3.**
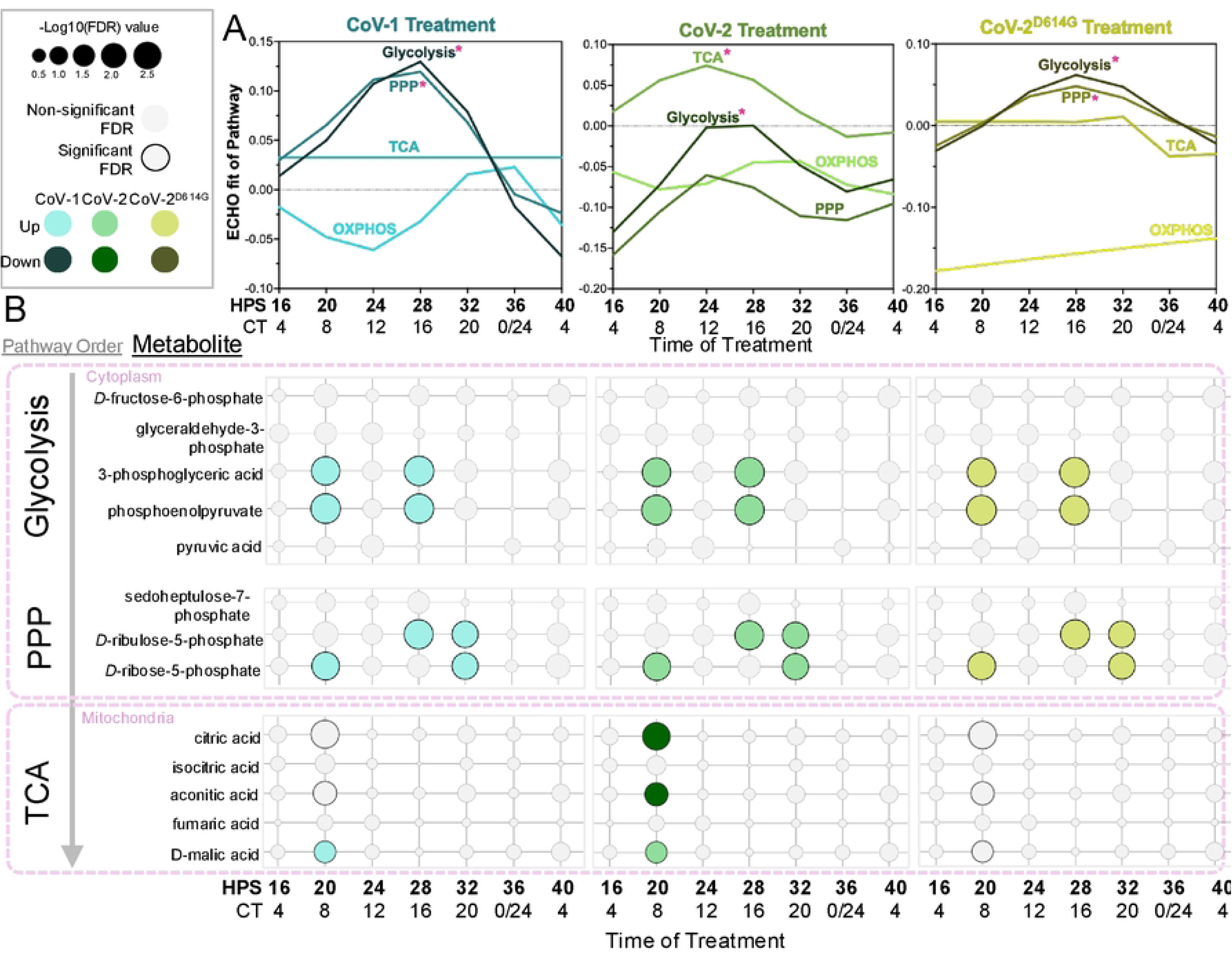
Metabolites in central metabolism are differentially regulated by spike protein exposure in a time-dependent manner. **(A)** ECHO fitted models of the log2FC of all proteins within the four central metabolism pathways in response to CoV-1, CoV-2, and CoV-2^D614G^ spike proteins at different times of the day (HPS/CT). Significant circadian fits are represented with an asterisk. **(B)** Bubble plot showing log2FC values of all detected central metabolite responses to spike proteins (left panel CoV-1, middle panel CoV-2, right panel CoV-2^D614G^, delineated by order in the pathway. Bubble size represents - log10(FDR), bubble color represents spike treatment, with lighter/darker shades representing significant up/downregulation, and outlined bubbles have a significant FDR < 0.05, as in the key.

Given the extensive time of day response of proteins in the metabolic pathways, we next extended our analysis to look at the differential temporal response of metabolites to spike proteins by applying a differential expression analysis to our metabolic data. We found that treatment with spike protein at HPS20/CT8 and HP28/CT16 gave the highest changes in central metabolite abundance, which parallels the increase in the differential responses noted in the glycolysis, TCA, and PPP pathway proteins (**Fig 3B**). DEMs included 3-phosphoglyceric acid and phosphoenolpyruvate in glycolysis, D-ribulose and ribose-5-phosphate from the PPP, and citric acid, aconitic acid, and malic acid from the TCA cycle (**Fig 3B**). Taken together, these data indicate exposure to spike proteins have a broad effect on metabolic pathways that could affect the immune response.

### Mitochondrial function and morphology show time-of-day response to SARS spike proteins

Mitochondria are closely tied to both central metabolism and immune function and, under non-stimulatory conditions, show rhythmic changes in fission and fusion morphologies that correspond to peaks in abundances of metabolites in the TCA cycle and OXPHOS proteins [8]. Moreover, changes to mitochondria energetics and morphology have been observed in other cell types following spike protein exposure [16–18,43]. Therefore, we next decided to measure if spike protein exposure affected changes to mitochondria in macrophages, if these changes were time-dependent, and if there were time-dependent changes in mitochondrial-associated lipids and metabolites.

We began by analyzing all significant DEPs for significant GO ontologies and noted significant enrichment for mitochondrial GO terms, particularly with CoV-2^D614G^ spike protein treatment **(S4A Fig)**. Given this enrichment, we tracked time-of-day effects of individual mitochondrial proteins to spike treatment, sub-setting all significant DEPs to categorize mitochondrial proteins according to their functional grouping using the MitoCarta3.0 database [44]. We found 47 mitochondrial DEPs in at least one treatment time point across all three spike protein treatments. Most proteins from the five main functional categories represented—OXPHOS, metabolite transport, lipid-related, morphology, and mitochondrial membrane potential (MMP)—were downregulated, including the cytochrome c oxidase complex, particularly at HPS20/CT8 **(Fig 4A, S4B)**. Knockdown of this complex in mouse macrophages is known to increase mitochondrial stress leading to increases in reactive oxygen species (ROS) and phagocytosis [45]. These data correlated with the downregulation of proteins in membrane potential **(Fig 4A)**. Additionally, many metabolite and lipid transport proteins were downregulated, including two key proteins (ADT1 and ADT2) within the ADP/ATP anti-port system which constitute the final step of OXPHOS [46]. Downregulation of these metabolite transporter proteins indicates that spike protein potentially uncouples important mitochondrial functions such as the TCA cycle and ATP/ADP anti-porting from downstream energy production [46]. CoV-2^D614G^ spike treatment at HPS20/CT8 also induced downregulation of a mitochondrial-related cardiolipin (CL(72:5)) while both CoV-2 and CoV-2^D614G^ spike protein treatment induced upregulation of an immune-signaling lysophosphatidylinositol lipid (PI(18:0/0:0)_A), demonstrating further disruptions to mitochondria and immune function on other levels of cellular physiology **(FIG 4B)** [47].

**Fig 4.**
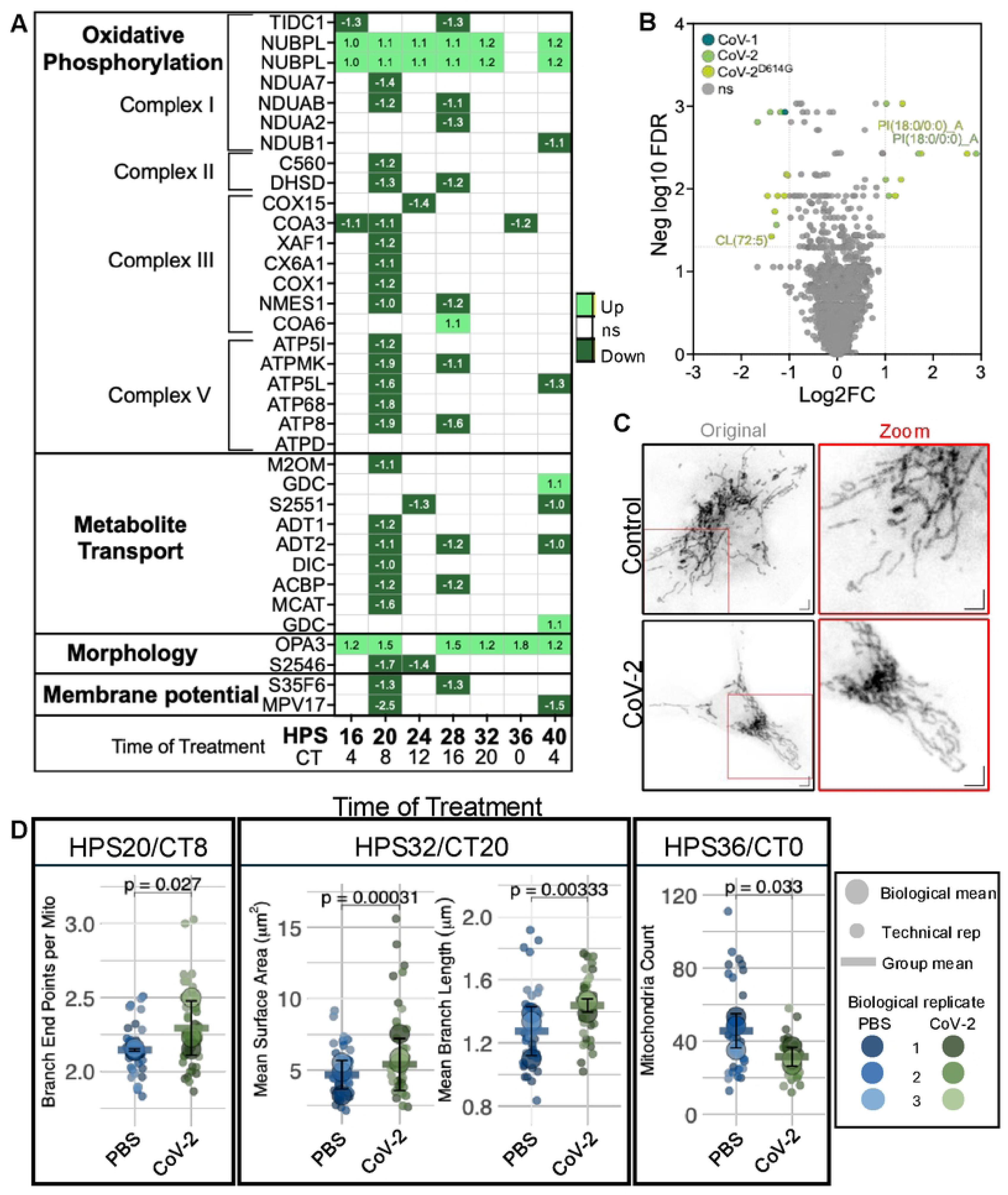
Mitochondrial function and morphology are altered in a time-dependent manner after spike protein exposure. **(A)** A chart of significantly up- or down-regulated log2FC values of mitochondrial proteins separated by functional category in response to CoV-2 spike protein (MitoCarta3.0 [44]). Internal numbers represent significant log2FC upregulation (≥1.0, light green) or downregulation (≤1.0, dark green). Timepoints shown are in HPS and CT. **(B)** Volcano plot showing differential level analysis of lipids responding to CoV-1, CoV-2, and CoV-2^D614G^ spike proteins at HPS20/CT8, colored by treatment if significant (ns = not significant, in gray). **(C)** Confocal maximum projections of representative fixed mBMDMs stained with MitoTracker^TM^ CMXRos Red following treatment with either PBS control or CoV-2 spike protein at HPS20. The original image (left) contains an ROI labeled in a red box, zoomed and represented to the right. Scale bar represents 2μM; images pseudo-colored using the inverted look-up table (LUT) in ImageJ/FIJI. **(D)** SuperPlots quantifying significant mitochondrial parameters at each time point. Biological replicates are represented by large, circled points; smaller points represent technical replicates; the line represents group mean; and error bars are ±1SD. Matching technical and biological replicates are represented by the same color saturation as indicated in the key. Statistics: linear mixed model, Holm-Šídák adjusted.

Given the downregulation of these pathways in response to spike protein exposure, we next measured if mitochondrial function was impaired. We first assayed mitochondrial membrane potential (MMP)—a readout of OXPHOS function—under non-stimulatory conditions to see if it followed a circadian pattern, as shown in another comparable study [48]. To do so, we synchronized our mBMDMs and incubated them with JC-10, a ratiometric dye that accumulates in mitochondria with high MMP. Readings of JC-10 ratios were recorded every four hours for 24 hours and modeled for circadian rhythms. We found that under non-stimulatory conditions, MMP followed a circadian rhythm, with low MMP levels coinciding with the highest percentage of fissioned mitochondria as seen in similar cell types by other groups (**S5A Fig**) [10,48]. Given this demonstration of circadian regulation of MMP, we next measured MMP in synchronized mBMDMs treated with spike proteins at the same 4-hr time points for 24 hrs as above. Using the ratio of MMP readouts of the spike-treated samples to vehicle-control treated samples, we noted rhythmic trends in the response to CoV-2^D614G^ spike protein, which was confirmed by ECHO **(S5B Fig)**. While no significant ECHO models were found for CoV-1 and CoV-2 treatment, we noted that they followed similar trends as CoV-2^D614G^ and displayed the lowest MMP levels at HPS20, which corresponded with the proteomic predictions (**S5B Fig**). When we compared MMP responses to classical pro- and anti-inflammatory stimuli—LPS and IL-4—we noted that the time-of-day trends in response to both stimuli were similar to spike protein treatment **(S5C Fig)**.

We next examined whether the weakly-rhythmic MMP responses to spike protein were concomitant with changes to mitochondrial morphology, which is known to impact OXPHOS function and is disrupted by spike protein exposure in other cell types [18,43,49,50]. In these experiments, we proceeded only with the CoV-2 spike protein, as this was the only variant to induce time-of-day changes to both proteins and metabolites within the TCA cycle. We circadianly synchronized mBMDMs and treated them with either CoV-2 spike protein, LPS, or IL-4 (the latter two of which are known to alter mitochondrial morphology via pro-inflammatory or anti-inflammatory pathways, respectively [51]) at four different time points (HPS20/CT8, HPS24/CT12, HPS32/CT20, and HPS36/CT0), targeting the treatment time points seen in the proteomics data when mitochondrial proteins were hypothesized to be affected. After a 6hr incubation with the respective treatments, we stained mitochondria with MitoTracker® CMXRos red dye, fixed, and imaged mitochondria from individual cells using fluorescent confocal microscopy **(Fig 4C, S5D)**. We used the FIJI plugin MitochondriaAnalyzer to threshold and analyze multiple mitochondrial morphology and network parameters [52]. Overall, treatment with CoV-2 spike protein induced changes to mitochondrial morphology that were time-dependent and mirrored the changes in both the proteome and membrane potential. Treatment at HPS20/CT8 significantly increased network branching in comparison to the vehicle control but had no effect on the other measured parameters **(Fig 4C, D).** Spike treatment at HPS32/CT20 caused increases to mean branch length and mean surface area, indicating mitochondria lengthening **(Fig 4D)**. Treatment at HPS36/CT0 led to decreased mitochondria counts and total branch end points **(Fig 4D)**. Mitochondrial responses to LPS and IL-4 were distinct from responses to CoV-2 spike protein in both treatment time and type of mitochondrial response **(S5D Fig** and **Supplemental Dataset)**. This indicated that spike protein induced a mitochondrial response that was influenced by the clock, but this response did not mimic the mitochondrial effects of the classical inflammatory response.

### Human macrophages mimic murine metabolic and immune-related circadian coordination

Given the time-of-day specific responses in mouse mBMDMs, we next characterized the response to SARS infections in human macrophages as, though mice can model the human immune response, human cell lines are a better exemplar [53]. To do so, we used human macrophages (hMΦ) derived from peripheral blood mononuclear cells (PBMCs) from an adult male human donor. These cells expressed macrophage-specific markers (CD14, CD11b/CR3, CD206), a macrophage differentiation marker (Mcl-1), and other lymphoid markers (CD4, CD2, CD19), confirming their identity as macrophages. We performed a similar clock synchronization protocol on the hMΦs as on the mBMDMs, whereby we grew them to 60-80% confluence, starved them in an FBS-free media for 24 hours, then subjected them to a 2-hour 50%-FBS serum shock before subjecting them to 2% heat-inactivated FBS supplemented media **(Fig 5A)**.

**Figure 5.**
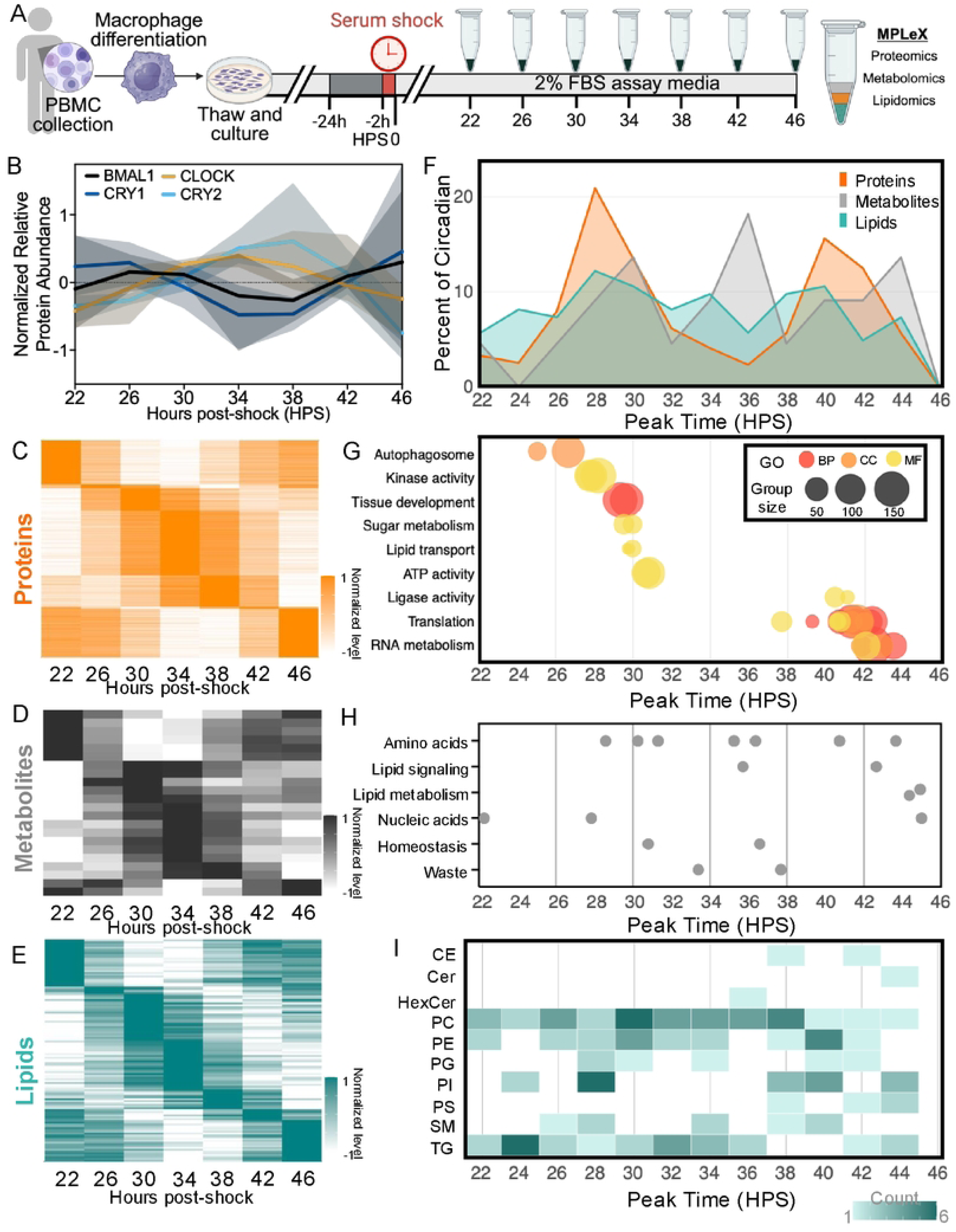
24-hr circadian oscillations in human macrophages time metabolic and immune processes across multiple omics scales. Schematic of the timeline for collecting multi-omics data from circadianly-synchronized human macrophages derived from peripheral blood monocyte cells (PBMCs). PBMCs derived into macrophages were cultured until 60-80% confluence, starved for 24hrs, then serum synchronized for 2hrs to reset the clock. Samples for proteomics, metabolomics, and lipidomics were collected every 4 hrs at hours post-shock (HPS) 22, 26, 30, 34, 38, 42, 46, and sent for mass spectrometry analysis using the MPLEx method. **(B)** ECHO model fit (line) of detected core clock proteins. Shaded portion represents ±1SD of the data. **(C-E)** Heatmaps showing ECHO fit models of relative high (dark) to low (light) abundance of circadian **(C)** proteins (orange), **(D)** metabolites (grey), and **(E)** lipids (teal) with significant circadian oscillations (period = 20-28 hrs, BH-adj p-val <0.05). **(F)** Overlay of peak phase distributions of circadian entities within each omics group by HPS. **(G)** Significant PSEA results grouped by functionally similar ontology terms. Each dot represents the mean peak phase time of an ontology term within its group. Ontology type is indicated by color (Biological Process (BP) = dark orange, Cellular Compartment (CC) = light orange, Molecular Function (MF) = yellow). **(H)** Circadian metabolites grouped by type according to KEGG annotations. Each dot represents peak phase time of one metabolite. **(I)** Heatmap showing peak phase time of circadian lipids within each class, binned by 2 hrs. Cholesteryl esters (CE), ceramides (Cer), hexosylceramides (HexCer), phosphatidylcholines (PC), phosphatidylethanolamines (PE), phosphatidylglycerols (PG), phosphatidylinositols (PI), phosphatidylserines (PS), sphingomyelins (SM), triglycerides (TG).

To begin, we first needed to characterize circadian oscillations in these cells, as little is known about circadian regulation in primary human macrophages. As with the mBMDMs, we harvested, pelleted, and flash froze the hMΦs at HPS22 and every subsequent 4 hours for the following 24 hours, giving us a total of 7 samples. These samples were then prepared and examined using relative quantification mass spectrometric analysis to identify global proteomics, metabolomics, and lipidomics using the MPLEx method (**Fig 5A**), and the resultant data were analyzed as it was for the mBMDMs.

Using the protein layer of the MPLEX, we identified a total of 85,276 unique peptides across 10,173 unique proteins. After imputing missing data and removing batch effects using LIMBR, and rolling the peptides up to the protein level, we noted 6,807 proteins that were reliably detected (>70% coverage) within our data set [24]. Using the metabolite layer of the MPLEX, we identified 114 metabolites, which were then subjected to LIMBR processing to impute missing data (if >70% coverage) and adjust for batch effects. This resulted in 107 metabolites which were reliably detected within our data set. Using the lipid layer of the MPLEX, we identified 437 lipids in both positive (247 lipids) and negative (190 lipids) runs. LIMBR analysis resulted in all 437 lipids being classified as reliably detected within our data set. PCA analyses of all three omics data sets, and correlation plots for the metabolomics and lipidomics data sets, indicated the three replicates were highly correlated (**S6A-E Fig**).

To identify circadian regulation within each data set, we used ECHO to model oscillations (damped, harmonic, forced) with periods ranging from 20-28 hours, as we empirically determined this to be the range of periods for the core clock proteins we detected (BMAL1, PER1, CRY1, and CRY2) (**Fig 5B**) [23]. Using these parameters, we identified 1,986 (29.2%) of proteins **(Fig 5C)**, 22 (20.8%) of metabolites **(Fig 5D)**, and 123 (28.1%) of lipids **(Fig 5E)** oscillated with a circadian period in our hMΦs. To visualize the coordination of circadian timing, we plotted the peak timing of all circadian proteins, metabolites, and lipids. We found patterns in the proteins from the hMΦs that were similar to our mBMDMs, where hMΦ proteins display a bimodal distribution twelve hours apart (HPS28 and HPS40). However, the metabolites and lipids had a broader range of peak timing (**Fig 5F**).

To determine which cellular processes were under circadian regulation, we employed phase-set enrichment (PSEA), which used Kuiper circular statistics and Gene Ontologies to determine statistical GO enrichment across the circadian day. PSEA uncovered translation, RNA metabolism, and RNA splicing peak together (HPS42), while immune-functions (tissue development, autophagosome), signaling (kinase activity), and energy/metabolism (sugar transferase, ATP activity, lipids, ligase activity) terms are enriched about 12 hours prior (HPS24-30), mirroring patterns observed in the analysis of mouse BMDMs from our previous work [10] (**Fig 5G**).

Given the consistent trend in metabolic regulation, we examined the circadian regulation of central metabolism in hMΦs. We found circadian control of key transition steps in glycolysis, the TCA cycle, and OXPHOS. These key transitions include the commitment of glucose to glycolysis **(S7A Fig)**, the transition from glycolysis to the TCA cycle **(S7A, B Fig)**, and the steps linking the TCA cycle and OXPHOS **(S7B, C Fig)**. Overall, the circadian regulation of the enzymes in central metabolism further mirrored the patterns observed in mice.

Although ECHO did not identify any central metabolites as circadian, other energy and immune-related metabolites were recognized as being under circadian control, including terms related to amino acids, nucleic acids, and lipid metabolism terms. Of note, nucleic acids peaked around HPS24, just prior to both the oscillating proteins involved in splicing and translation events **(Fig 5H)**. In parallel with circadian metabolites involved in immune regulation, lipid signaling metabolites involved in immune regulation (myo-inositol and scyllo-inositol phosphate, the latter being protective against toxic forms of amyloid-beta) were also found to be under circadian control (**Fig 5H**) [54,55].

In our lipidomic data, we found oscillating lipids that were coordinated with immune signaling [56]. We noted that, while nearly all detected classes of lipids were circadianly regulated, the classes phosphatidylinositols (PI), triglycerides (TG), and sphingomyelins (SM) held the highest percentage of circadian entities **(S7D Fig)**. PI lipids peaked twice (HPS28, HPS40), and these peaks occurred around the same time of day as their head group, myo-inositol **(Fig 5E, I)**. Moreover, PI lipids are activated as signaling molecules through kinase phosphorylation, and we noted similar peak timing in a large subset of PI lipids at the same time as kinase activity terms (HPS28) **(Fig 5G and I)** [57]. Finally, three PIs directly linked to immune function—two lysophosphatidylinositols (PI[18:1/0:0_A], PI[20:3/0:0_A]) and one lipokine (PI[18:1/18:1]) were circadianly regulated **(S7E Fig)**.

Given the tie between the immune response and mitochondria, and the regulation of mitochondria in the mBMDMs, we next filtered our circadian dataset for mitochondrial proteins using the MitoCarta3.0 database [10,44,58]. After separating the circadian mitochondrial proteins into functional groupings, we noted that a high proportion were related to the mitochondrial central dogma, particularly ribosomal subunits (translation) and DNA and RNA binding proteins **(S7F Fig)**. Additionally, several key mitochondrial morphology regulating proteins (including MFN2 (fusion), DNM1L and AKAP1 (fission)) were observed to oscillate, suggesting similar mitochondrial morphological gating as we noted in mBMDMs previously **(S7F Fig)** [10,59,60].

In total, across all three omics levels, we found the signatures of circadian regulation on energy regulation, immune signaling factors, protein translation, and mitochondrial dynamics in human macrophages that mirrored what was seen in mouse macrophages [10]. While circadian regulation of specific proteins, metabolites, or lipids may not necessarily be the same between both species, this conservation of function appears to preserve circadian temporal regulation within major macrophage cellular functions between murine and human cells.

### Exposure to spike proteins affects central metabolic pathways in human macrophages

Having analyzed the circadian oscillations in the human macrophages and found many overlaps with the murine macrophages, we hypothesized the hMΦs would have similar time-of-day responses upon exposure to spike protein. For these experiments, we chose the WT CoV-2 and CoV-2α spike protein variants as the latter was, at the time of these experiments, the current variant of concern that displayed increased transmissibility over WT, although it still retained ∼98% similarity **(Fig. S2A)** [61]. To investigate the impact of the time of day on the immunometabolic response, we exposed human-derived circadianly-synchronized macrophages to the S1 spike protein of SARS-CoV-2 and CoV-2α at seven different time points across the circadian day (HPS16, 20, 24, 28, 32, 36, 40) for 6 hrs, in triplicate **(Fig 6A)**. Cell pellet samples from these time points were extracted and analyzed for proteomics, metabolomics, and lipidomics using the MPLEx method as described above **(Fig 6A)**. We cleaned and pre-processed the data sets using LIMBR, and PCA and correlation plots indicated high similarity between replicates and tight clustering of treatments **(S6A, C-F Fig)** [24]. We then analyzed each omics set for differential levels relative to the vehicle control using the edgeR package in R as described above using a higher log2FC cutoff (|log2FC| ≥1.5) to increase stringency, as our human data displayed increased levels of DEPs as compared to the mouse data [35].

**Figure 6.**
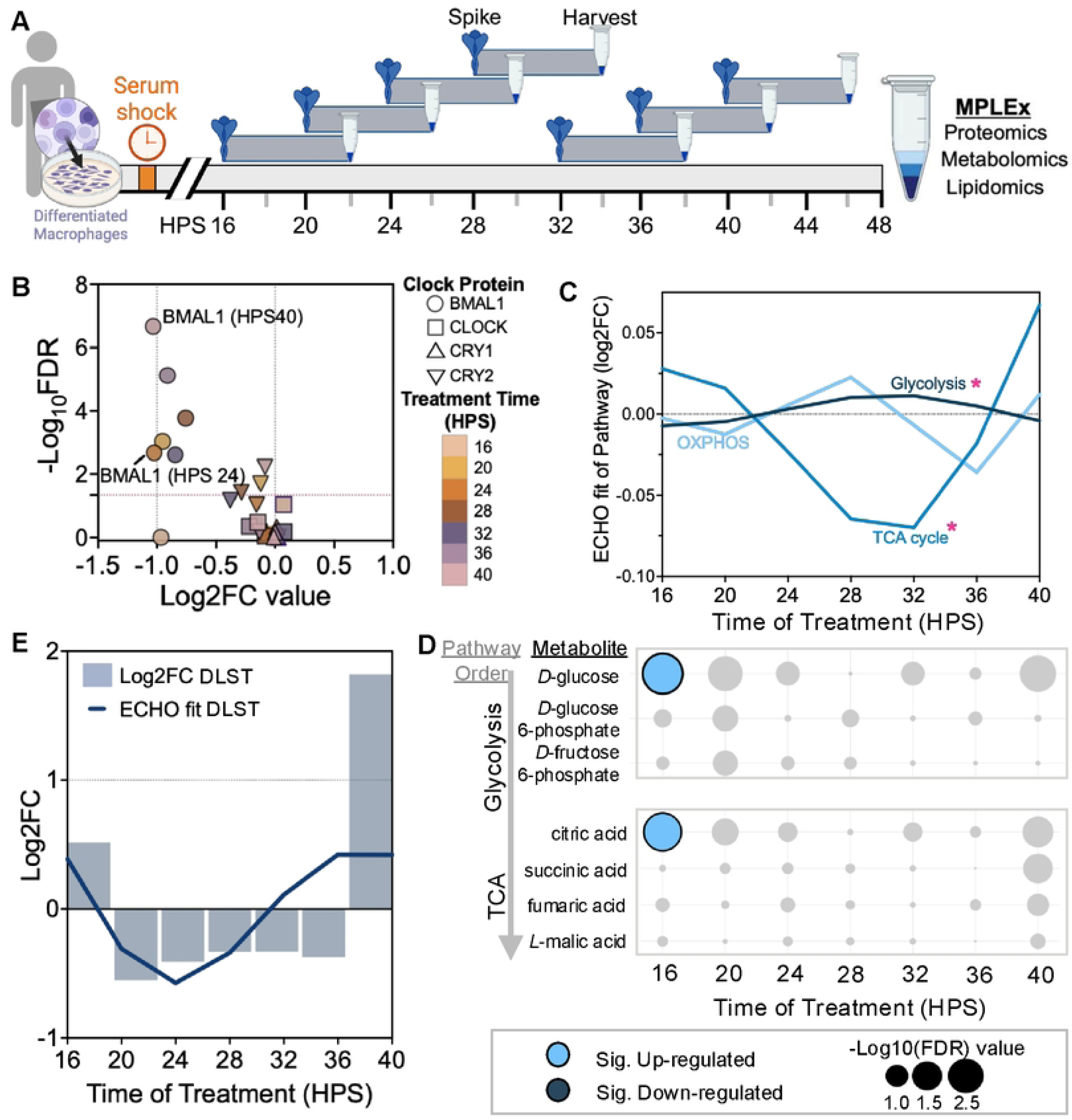
Clock proteins and metabolic pathways in human macrophages respond in a time-of-day dependent manner to CoV-2 spike protein. **(A)** Diagram showing the experimental setup for exposing circadianly-synchronized human macrophages to SARS-CoV-2 spike protein and harvesting cell pellets for mass spectrometric analysis using the MPLEx method. **(B)** Volcano plot showing differential levels of circadian core clock proteins at each treatment time point. Shape indicates clock protein; color denotes the time of spike protein treatment. Dotted lines indicate significant log2FC and -log10(FDR) cutoff values. **(C)** ECHO model fit lines to the log2FC values of all central metabolic proteins, separated by pathway, in response to CoV-2 spike. Significant fits are denoted by (*). **(D)** Bubble plot of all detected metabolites within central metabolic pathways glycolysis and TCA cycle. Color indicates significant up (light blue) or down (dark blue) differential levels, while bubble size denotes –log10 FDR value. Metabolites are listed in the order they follow in each pathway. **(E)** ECHO fit (line) and corresponding log2FC (bar) of circadian response proteins DLST at each treatment time point.

Contrary to what we observed in mice, human macrophages showed no overall variation in the number of DEPs, with ∼375 DEPs proteins at all time points except HPS40 **(S8A, B Fig)**. DEPs from all time points roughly shared equal proportions of up- and down-regulated proteins for both CoV-2 and CoV2α **(S8A, B Fig)**. In contrast to the mBMDMs, we did observe downregulation of BMAL1 at both HPS24 and HPS40 treatment times with CoV-2 (but not CoV-2α), suggesting spike exposure in hMΦs may affect some aspects of the core clock with certain spike variants **(Fig 6B, S8C).**

In keeping with the trend of circadian metabolic regulation, we next analyzed the response of central metabolism proteins and metabolites to spike protein in hMΦs. We again used ECHO to model the log2FC of all proteins over time within glycolysis, TCA cycle, and OXPHOS to identify circadian rhythms in their responses [23]. We found both CoV-2 and CoV2α treatments induced significant oscillatory responses in glycolysis and the TCA cycle in an anti-phase manner **(Fig 6C, S8D, F Fig)**. In fact, the trough in glycolysis (∼HPS16) corresponded with a significant increase in the metabolite *D*-glucose, while the peak in TCA cycle response corresponded with a significant increase in citric acid **(Fig 6C** and **E, S8D)**. When we applied our significant circadian response analysis approach to individual proteins (|log2FC| ≥1) **(Fig 2D)**, we noted that the response of TCA protein Dihydrolipoyllysine-residue succinyltransferase component of 2-oxoglutarate dehydrogenase complex (DLST) also followed a circadian oscillation **(Fig 6E)**. Overall, these central metabolic responses closely resembled those observed in mBMDMs, indicating conservation of spike protein response between the two species. These data suggest the TCA cycle as a key responder to spike protein in circulating monocytes in both human and mouse.

To determine if the circadian response effect of spike proteins also affected mitochondria in hMΦs, we filtered the list of proteins with a significant circadian response (CRPs) for mitochondrial proteins. Of the five CRPs we identified, two of these, PHB2 and TIM14, which oscillate concordantly, preserve mitochondrial membrane integrity through cardiolipin regulation **(S8G Fig)** [62,63]. In addition, we also identified the protein SOD2, a primary scavenger of reactive oxygen species (ROS) in mitochondria that has been shown to respond to spike proteins **(S8G Fig)** [43,64,65]. Finally, we found several proteins responsible for macropinocytosis, a key pathway stimulated after spike protein exposure which increases spike uptake [66]. We noted that one of the main activators of macropinocytosis, PKCδ, oscillated in response to CoV-2 spike protein exposure **(S8H Fig** [67,66]. In total, these data highlight a conservation of the circadian mitochondrial and metabolic response to spike protein between humans and mice.

## Materials and Methods

### Mouse and Human macrophage sources, culture, and circadian synchronization

PER2::LUC C57BL/6J male mice aged 3–6 months sourced from The Jackson Laboratory (Strain # #:006852) and bred in-house were used for all experiments involving primary BMDMs[68]. Mice were housed under controlled temperature and humidity conditions and maintained on a strict 12 h light:12 h dark cycle. They were provided standard rodent chow and water ad libitum. Euthanasia was performed using CO₂ inhalation (50% chamber volume per minute) followed by cervical dislocation. All animal procedures were conducted in compliance with the National Institutes of Health Office of Intramural Research guidelines and were approved and supervised by the Rensselaer Polytechnic Institute Institutional Animal Care and Use Committee. Monocytes were derived from bone marrow collected from the fibulas and tibias of PER2::LUC C57/BL6 male mice, aged 3-6 months, and cultured at 37°C and 5% CO_2_ as described by Collins [10]. Briefly, 1×10^6^ monocytes were plated on 35mm dishes and differentiated into macrophages in DMEM media supplemented with 10% FBS and 0.1μg/mL M-CSF (Prospec Cat# CYT-439) (media replaced every 3 days). After 7 days of differentiation, the resultant macrophages were washed with PBS, starved for 24hrs in Leibovitz media (Gibco™, ThermoFisher Cat#21083027) with M-CSF but devoid of FBS, and then shocked for 2hrs in Leibovitz media with M-CSF and 50% FBS to synchronize circadian rhythms via the serum response element on the PER2 promoter [69]. The synchronized macrophages were then given Leibovitz media supplemented with 10% FBS and luciferin (GoldBio Cat# LUCK-100) and luminescence was tracked in a LumiCycle32 (Actimetrics). All experiments were performed in independent biological triplicate (e.g. from three male mice) unless otherwise indicated.

Human macrophages were sourced from CelProgen (Cat# 36070-01) from a 35-year-old white male (Donor #30, Lot#1414270) and were originally collected in 2018, prior to the COVID-19 pandemic. The frozen macrophages were reanimated and cultured in human macrophage serum-free media (CelProgen Cat#M36070-01) supplemented with 10% heat-inactivated FBS (HI-FBS), and passaged 1:3 every 2-3 days, or when confluent. For circadian synchronization, 5×10^4^ cells were plated into an uncoated 35mm TC dish. When the cells reached 60-70% confluence, their media was changed to Leibovitz media devoid of FBS for 24hrs (starve media). Circadian synchronization was achieved following a 2hr incubation with 50% HI-FBS, and cells were changed into Leibovitz media with 2% HI-FBS. All experiments were performed in technical triplicate unless otherwise indicated.

### Time course, spike protein exposure, and cell collection for multi-omics

Starting at 16hrs post-50% FBS shock (HPS), mouse BMDMs or human macrophages were exposed to 11.2nM (1μg/mL) of SARS spike protein in PBS as indicated every 4 hours for 24 hours, for a total of 7 time points (at HPS16, 20, 24, 28, 32, 36, and 40). mBMDMs were incubated for 6 hours with either PBS (vehicle control), SARS-1 spike protein, SARS-COV-2 S1 spike protein, or SARS-COV-2^D614G^ S1 spike protein (SinoBiological Cat#40150-V08B1, 40591-V08B1, 40591-V08H3, respectively) as indicated. Human macrophages were incubated with either PBS (vehicle control), SARS-COV-2 S1 spike protein, or SARS-COV-2α S1 spike protein as indicated (SinoBiological Cat# 40591-V08B1 and 40591-V08H12, respectively). Six hours after the addition of spike protein, mBMDM or human macrophage samples were washed with warm PBS, harvested using CellStripper (Corning^TM^ Cat# 25056CI), pelleted, flash frozen, and sent to the Pacific Northwest National Lab (PNNL) for mass spec analysis to measure proteomic, metabolomic, and lipidomic levels using the MPLEx method [22]. The program MOE (Molecular Operating Environment, 2024.06) was used to perform a sequence alignment to compare the S1 spike protein amino acids of the SARS-CoV variants [70].

### Proteomics

Protein preparation and mass spectrometry for both mouse and human macrophage samples were all completed at PNNL following the MPLEx method [22]. Briefly, cell pellets were digested with trypsin and labeled with one of 16 unique tandem-mass-tag labels. Labeled peptides were separated using off-line high pH (pH10) reversed-phase separation with a Waters XBridge C18 column using an Agilent 1200 HPLC system coupled to a Q Exactive Hybrid Quadrupole Orbitrap Mass Spectrometer operated with an NSI ion source (positive spray voltage = 2200V, ion transfer tube temperature = 250°, default charge state = 2, run time = 120 min). The parent MS scan was obtained with the following settings: 120,000 resolution, AGC target = 400,000, RF lens = 30, scan range of 300-1800 m/z, inclusion of charge states 2-6, dynamic exclusion settings of 45s duration, mass tolerance = 10, maximum intensity = 1e+20, minimum intensity = 50,000, and the IntensityThreshold filter. For MSn Scans, HCD was performed on isolated parent ions isolated with an isolation window = 0.7. Normalized collision energy was used at 32% with a maximum injection time of 86 ms. The dynamic exclusion time was set to 45s. The resolution for MSn scans was set to 50,000, and the AGC Target = 125,000 with normalized AGC target = 250.

The resultant MS/MS spectra from mouse samples were searched and mapped against M_musculus_UniProt_SPROT_2019-10-11 and a database for common human and bovine contaminants. Resulting MS/MS spectra from human samples were searched and mapped against H_sapiens_UniProt_SPROT_2021-06-20, SARS-CoV-2_Covid-19_UniProt_2020-10-22, and Tryp_Pig_Bov. Measured mass accuracy and MSGF spectra probability were used to filter identified peptides to a <0.4% false discovery rate (FDR) at spectrum level and <1% FDR or <5% FDR at the peptide level, using the decoy approach. TMT reporter ions were extracted using the MASIC software with a 20ppm mass tolerance for each expected TMT reporter ion, as determined from each MS/MS spectrum [71]. Only peptides mapped to a unique protein were used for subsequent analysis.

Peptide counts were log2-normalized and then run through LIMBR (v0.2.10), an SVA method designed to impute missing values and model and remove batch effects within circadian mass spec data in a python environment [24]. Note that the non-treated control human macrophage samples were additionally normalized to the pooled reference samples for each batch before log2 normalization and running LIMBR. Within LIMBR, the default KNN-based method (n = 10) was used to impute missing values for entities that had >70% data coverage across all time points; entities with <70% coverage were removed. Next, all mouse control and spike-treated samples, along with the human spike-treated samples, were modeled for batch effects in LIMBR’s circadian mode using the default settings, including 100 permutations. Post-LIMBR peptide abundances were rolled up to the protein level by taking the mean value of all peptides assigned to a given protein.

### Metabolomics

Metabolites for both mouse and human macrophage samples were all completed at PNNL following the MPLEx method and further sent for analysis using GC-MS as previously described [22,72]. Briefly, the metabolite extract layer derived from the MPLEx method was injected into an Agilent 7890A gas chromatograph coupled with an Agilent single quadrupole 5957C mass spectrometer. *m/z* spectra were obtained and processed through MetaboliteDetector and in-house curated libraries based on an augmented version of the Agilent Fiehn Metabolomics Retention Time Locked (RTL) Library [73,74]. ^13^C-labeled fumaric acid was used as an internal standard control for GC-MS and was included within the LIMBR imputation and batch effect steps below. Metabolite counts for both mouse and human samples were log2-normalized and run through LIMBR, as described above for the protein peptides.

### Lipidomics

Lipids for both mouse and human macrophage samples were extracted from the metabolite layer derived from the MPLEx method and further sent through LC-ESI-MS/MS analysis as previously described [22,75]. Briefly, lipid extracts were injected into a Waters Aquity UPLC H class system which was coupled with a Velos-ETD Orbitrap mass spectrometer. Samples were analyzed in both positive and negative ionization modes using higher-energy collision dissociation (HCD) and collision-induced dissociation (CID). The resulting MS/MS spectra were analyzed using LIQUID to determine lipid identity and lipid nomenclature was assigned based on LipidMaps [75,76]. Lipid abundances from each ionization mode for both mouse and human samples were log2-normalized and run through LIMBR as described above for the protein peptides and metabolites, except for imputation, as there were no missing values.

### Data Validation

Both PCA analyses and correlation plots were used to validate all omics data for variation between samples, replicates, and time points. Principal Components were calculated using the prcomp() function in base R, and Pearson’s Correlation was performed using the cor() function in base R to determine correlation values (R v4.2.2 [77]).

### Circadian Detection

Post-LIMBR protein, metabolite, or lipid abundances from the vehicle control data sets were free-run through ECHO version 4.0 [23] with options selected for data smoothing, normalization (z-scoring), and linear detrending. To identify circadian entities, proteins, metabolites, and lipids were only considered circadian if they had a best-fit model of damped, harmonic, or forced, with a period of 20-28 hours and a BH-adj p-val <0.05. To convert our time course in HPS to CT, we first adjusted all protein, metabolite, and lipid circadian oscillations from ECHO to a standard period of 24hrs using the equation: 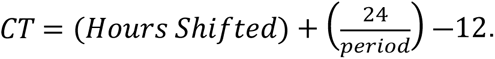 Next, we aligned our *in vitro* PER2 oscillations in our proteomics data to previous *in vivo* PER2 oscillations to infer CT such that HPS16 = CT4 [10,11].

### Differential levels of proteins, metabolites, and lipids

Differential levels of un-logged proteomics, metabolomics, or lipidomics data was analyzed using the Bioconductor package edgeR [35] (version 4.0.5) in R (Version 4.2.2). Each omics set and each time point was run through this analysis separately in edgeR’s multivariate mode which used a one-way ANOVA-like test to make comparisons between control- and spike-treated proteins and determine log2FC value and accompanying statistical F-test for each. For mouse, the significance thresholds were set at |log2FC| ≥1.0 and FDR <0.05; while for human, the significance thresholds were set at |log2FC| ≥1.5 and FDR <0.05. Upregulation was defined as ≥1.0 (mouse) and ≥1.5 (human), and downregulation as ≤ -1.0 (mouse) and ≤ - 1.5 (human).

### Protein and Metabolite GO analyses

KEGG database (Release 109.0, January 1, 2024) was used to identify proteins within central metabolic pathways for both circadian proteins and differentially-expressed proteins, as indicated [78]. Lists of central metabolic pathway proteins and metabolites were retrieved from KEGG (Glycolysis/Gluconeogenesis (*Mus musculus)*: mmu00010, Pentose Phosphate Pathway (*Mus musculus)*: mmu0030, Citrate Cycle (TCA Cycle) (*Mus musculus)*: mmu00020, Oxidative Phosphorylation (*Mus musculus)*: mmu00190, Glycolysis/Gluconeogenesis (*H. sapiens)*: hsa00010, Pentose Phosphate Pathway (*H. sapiens)*: hsa0030, Citrate Cycle (TCA Cycle) (*H. sapiens)*: hsa00020, Oxidative Phosphorylation (*H. sapiens)*: hsa00190) [25,78]. Where indicated, metabolites were categorized using KEGG Brite “Compounds with Biological Roles” tool. Version Sept. 18, 2022. StringDB (v12.0) was used to quantify protein relationships and ontologies of the connected significant circadian response proteins (CRPs) [36]. PantherGO (v17.0) statistical overrepresentation test was used to determine significant pathways within the significant differentially-expressed proteins. The query proteins were compared against a background of all reliably-detected proteins within the proteomics data set [79]. MitoCarta database (v3.0) was used to find and assign functional categories to mitochondrial proteins within the significantly differentially-expressed protein set for both human and mouse [44].

### Long-term tracking of circadian rhythms in response to spike protein via PER2::LUC

mBMDMs were collected, differentiated, and circadianly synchronized, and given Leibovitz media supplemented with 10% FBS and 250μM luciferin (GoldBio Cat# LUCK0100) and spike protein was added as described above. Bioluminescent traces were measured on a Lumicycle32 (Actimetrics) for up to 5 days following spike protein exposure. To prepare for ECHO analysis, counts/second for each biological replicate were binned into two-hour blocks, and the average count/second was used to represent the biological replicate at that time point. The starting point for assessing treatment effects was deemed to be the start of the treatment, and the end point was 72 hours following. Each time point was modeled separately in ECHO (v4.0) due to the different start and end times, and each biological replicate was treated separately to perform statistical analyses [23]. Rhythms were considered circadian if they had a best-fit model of damped, harmonic, or forced, with a period of 20-28hrs and a BH-adj p-val <0.05. A RM two-way ANOVA with Geisser-Greenhouse correction and post-hoc Dunnett’s multiple comparison test was used to determine statistical significance in treatment conditions at each time point compared to the control (GraphPad Prism Version 10.5.0 for Mac OS).

### Cytokine panel analysis

mBMDMs were collected, differentiated, circadianly synchronized, and treated with spike proteins every four hours beginning at HPS16 as described above. Following six hours of incubation with the spike protein, LPS (0.5μg/mL, Invitrogen Cat# 00-4976), or IL-4 (1ng/mL, Biotechne Cat# 404-ML/CF), the resultant macrophage supernatant was collected, centrifuged to remove cells and debris, and analyzed using a sandwich ELISA approach for the cytokines IL-6, TNF-alpha, IL-10, and TGF-beta-1. Cytokine concentrations were quantified using the corresponding DuoSet ELISA kits (R&D Systems, Cat# DY406, DY410, DY417, DY1679) in 96-well plates according to the manufacturer’s protocols (R&D Systems). The corresponding macrophages were measured for viability to normalize the cytokine concentrations using trypan blue (0.4% BioRad Cat#1450021). All experiments were run in biological triplicate (3 mice) and technical duplicate (i.e., the same supernatant sample was divided between two ELISA wells). Absorbances were read at 450nm with wavelength correction read at 570nm on a Tecan i-control infinite 200 Pro version 2.0.10.0. GraphPad Prism (Version 10.5.0 for Mac OS) was used to interpolate concentrations from a 7-point sigmoidal standard curve using kit-provided standards. The mean value of each technical duplicate was used to represent each biological replicate value. Resulting cytokine concentration values were normalized to the total number of live cells. GraphpadPrism (Version 10.5.0 for Mac OS) was used to perform an RM two-way ANOVA with post-hoc Dunnett’s multiple comparison test to determine statistical significance in treatment conditions at each time point compared to the control.

### Mitochondrial Membrane Potential

mBMDMs from a total of four male mice were collected, seeded in technical quadruplicate in 96-well plates at 2.5×10^5^ cells per well, differentiated, circadianly synchronized, changed into 100μL assay media (10% FBS, DMEM, no luciferin), and treated with spike proteins every four hours at HPS16, 20, 24, 28, 32, 36, and 40 as described above. Following six hours of incubation at each time point with either spike protein, LPS (0.5μg/mL, Invitrogen Cat# 00-4976), or IL-4 (1ng/mL, Biotechne Cat# 404-ML/CF), mitochondria membrane potential was measured using the JC-10 kit from Abcam (Cat#ab112134) according to the manufacturer’s directions. In brief, mBMDMs were incubated with JC-10 dye for 40 min, followed by addition of Buffer B. Fluorescent excitation/emission readings were immediately taken at Ex/Em = 490/525 (cutoff at 515nm, gain = 150); and Ex/Em 540/590 (cutoff at 570nm, gain = 180) on a Tecan i-control infinite 200 Pro version 2.0.10.0 plate reader. Values from each channel were normalized to the average value of those respective channels from 3 wells that contained unstained cells in assay media. The ratio of 590/525 channels was calculated and subjected to outlier analysis using the modified z-score method in R, and identified outliers were removed. These technical replicates were then averaged to and used to denote the biological replicate value. To identify circadian rhythms in the spike protein-treated data, 590/525 ratios of spike treated samples were taken as a percentage of the vehicle control and modeled for rhythms in ECHO (v4.0) [23]. Rhythms were considered circadian if they had a best-fit model of damped, harmonic, or forced, with a period of 20-28hrs and a BH-adj p-val <0.05.

### Mitochondria Morphology Experiments

2.5×10^5^ mBMDMs were seeded onto sterile 1.5 poly-L-lysine-coated (Sigma-Aldrich Cat#P4707) glass coverslips in 35mm cell culture dishes, grown, and circadianly synchronized as described above. Treatments were added at either HPS20, 24, 32, or 36, as indicated. 15 minutes prior to the end of each time point, 100nM MitoTracker™ Red CMXRos (ThermoFisher, Cat#M7512) was added to the cell culture media and incubated at 37°C for 15min. Cells were washed 3x with warm PBS, followed by fixation with 4% PFA/PBS at room temperature for 15 minutes. Cells were washed in RT PBS for 5 min 3x, cleared with 100% methanol for 15min, washed in RT PBS 3x, and left to dry in the dark. Once dry, the coverslips were mounted onto glass slides with mounting media (Sigma-Aldrich, Cat#06522), let dry in the dark overnight at 4°C, and sealed with clear nail polish the following day.

### Imaging

Mitochondria were imaged on a Leica SP8 confocal system with 100x oil objective (1.40NA) and white light laser set to 70% power. Images of 10-14 randomly selected mBMDMs were imaged for each time point, each treatment, and each biological replicate. Z-stack images spanning the whole height of each cell were taken at a Nyquist sampling distance of 0.30μM. To detect stained mitochondria, excitation was set to 578nm and emission to 584-657nm. Images were taken in a 1024 x 1024 format, 200 speed, line average of 4, with pinhole set to 1.00AU. Signals were detected using the built-in HyD detector with the smart gain optimally set for each image.

### Image Analysis

Z-stacks of each cell were cropped for ROI, randomized, and blinded for analysis in FIJI/ImageJ (version 2.16.0/1.54p) using the Mitochondria Analyzer plug-in (V2.3.1) [52]. Within this plug-in, 3D images were adaptively thresholded using a weighted mean for C-value and Block size, despeckled, and outlier-removed. Resulting thresholded images were then measured within the same package for all mitochondrial morphology parameters. Each mitochondrial parameter was analyzed separately for outlier detection using the interquartile range (IQR) method in R. The mean value of the resulting outlier-removed technical replicate values was used to represent the biological value. The linear mixed model was used to detect statistically-significant changes between the control and treated (CoV-2 spike protein, LPS, or IL-4) samples at each time point using the R-based packages emmeans, lmertest, and lme4 [80–82].

### PSEA Analysis on Human Macrophage Circadian Proteins

Phase-set enrichment analysis (PSEA, v1.1) was conducted on the circadian proteins from the untreated human samples using the java-based program PSEA to assess coordinated circadian regulation across functional pathways [83]. To allow for comparison of phase differences between circadian proteins, the peak timing for each circadian protein—as reported by ECHO—was adjusted to a 24-hour period using the equation: *Hours shifted*(24/period)*. These adjusted peak phase values were used as input into the java-based app and protein sets were defined using Hallmark Gene sets and Gene Ontology (GO) biological process (BP), cellular compartment (CC), and molecular function (MF) annotations from MSigDB (Human MSigDB v2025.1.Hs updated June 2025) [84–86]. Enrichment was calculated using circular Kuiper statistics to test whether proteins within a set shared phase clustering compared to the background circadian proteome. All parameters were set to the defaults for minimum gene sets and maximum simulations number in the java app. Results with q-values < 0.05 (vs. background) were considered significant, and mean phase values of functionally similar significant ontology terms were adjusted to fall within the experimental timeframe, then consolidated into manually-curated categories.

## Discussion

Innate immune cells, particularly macrophages, are heavily controlled by the 24-hour rhythms of the circadian clock, especially in immunometabolic pathways. However, little work has examined how time-of-day influences the macrophage response to stimuli at the molecular level. Here, we bridged this gap to show that the circadian clock plays a role in controlling the immunometabolic response across molecular levels and species. We were able to demonstrate that not only were immunometabolic protein pathways timed by the clock, but that the resultant metabolites and lipids also fell under circadian control in both mice and humans. Moreover, we showed that the clock times the response to immune stimuli (exposure to SARS spike proteins) in a similar manner across species, demonstrating the importance of circadian timing in response to pathogens **(Fig 7)**. Our work demonstrated that central metabolism and mitochondria, but not the overall core clock or immune pathways, responded differentially depending on the time of day of exposure.

**Figure 7.**
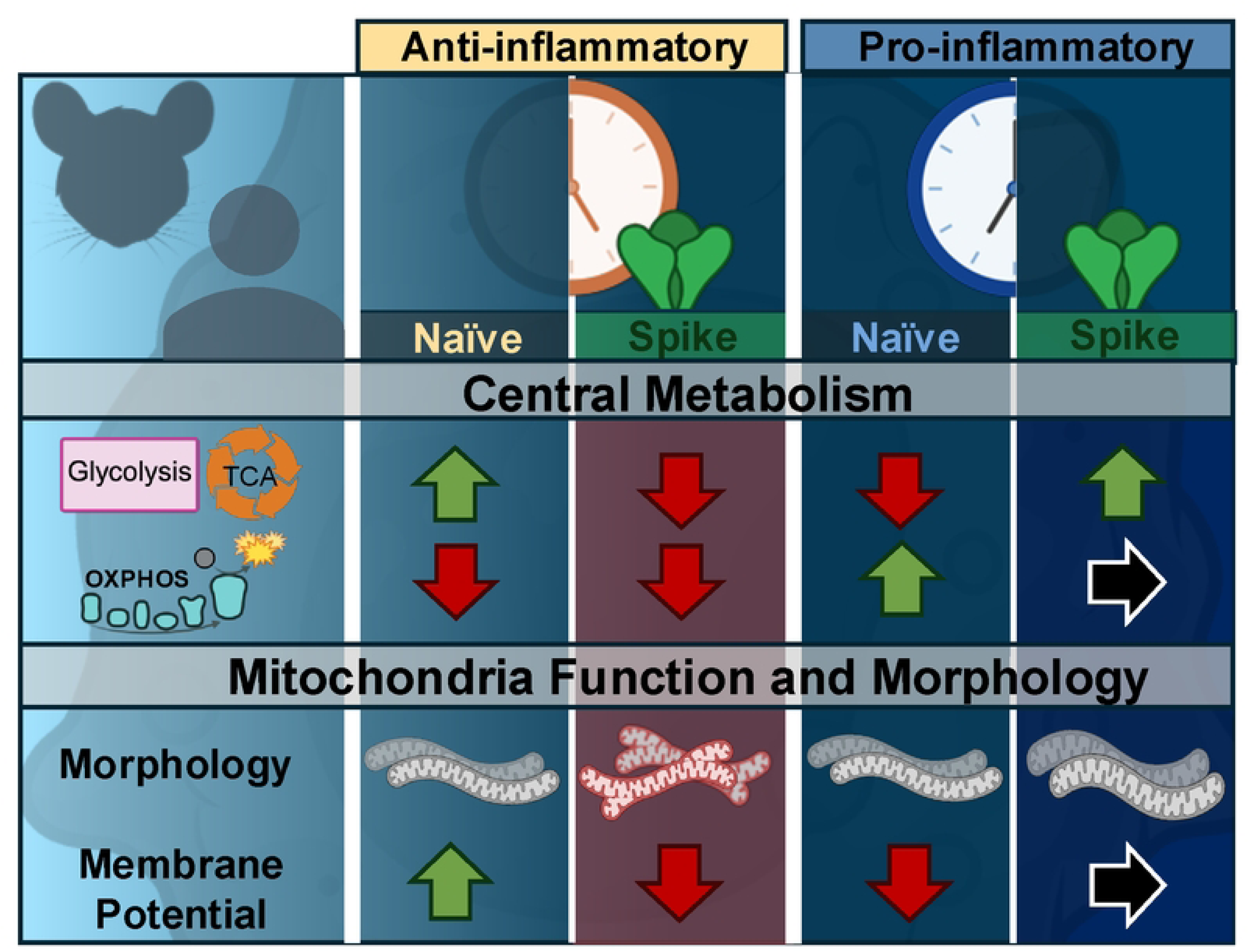
Spike proteins disrupt macrophage immunometabolism in a time-dependent manner. SARS-CoV-derived spike proteins elicited temporally-specific macrophage responses in both central metabolic function and mitochondrial function/morphology. During the circadian anti-inflammatory phase, when the proteins and metabolites in glycolysis, the TCA cycle, and MMP are high (green arrow), and OXPHOS is low (red arrow), exposure to the SARS-CoV spike protein treatment decreased protein levels and metabolites involved in central metabolism, decreased MMP pathways (red arrows), and increase mitochondrial branching. During the circadian pro-inflammatory phase, when proteins and metabolites in glycolysis, the TCA cycle, and MMP processes are low (red arrow), and OXPHOS is high (green arrow), exposure to the SARS-CoV spike protein either increased (green arrow) or maintained (black arrow) these processes and increased mitochondrial surface area.

Upon stimulation by a pathogen, the classic model of the immune response is that macrophages can rapidly polarize into a pro-inflammatory state by activating inflammatory immune pathways [2]. This polarization alters the rhythmicity of the circadian clock by changing the period and amplitude of the rhythm of the core clock proteins [37]. We confirmed this response in mouse macrophages using LPS treatment, noting increased cytokine secretion **(Figure S3F, G)** and alteration of core clock phase and amplitude **(Figure S2E)**. Surprisingly, our data showed no changes to the core clock proteins upon exposure to spike protein and no evidence of a temporal response in classical immune pathways within the proteomics data in mice **(Figure 2, 5, S2B-D, S3A-D)**. While moderate increases in both IL-6 and TNFα were noted, these trends were not specific to the time of day of exposure. Taken together, these data imply that although the spike protein variants stimulate modest inflammatory output, there is little evidence for temporal responses within classical inflammatory pathways, including inflammatory proteins or cytokines.

Observing no temporal responses in classical immune polarization in mBMDMs, we next turned to immunometabolic responses to see if spike proteins were altering energy regulation as seen in response to spike proteins in other cell types [17,43]. Classical pro-inflammatory stimuli alter central metabolism by disrupting the TCA cycle and OXPHOS to enhance fast energy generation via glycolysis, increasing the abundances of metabolites upstream of inflammatory pathways [2]. Here, we observed similar changes within central metabolic pathways in response to spike proteins, which were also time of day dependent. BMDMs treated around CT8, when glycolysis and TCA cycle pathways are at their zenith and OXPHOS abundance is at its nadir, saw a decrease in the abundances of glycolysis proteins and increase in the abundance of end-stage glycolysis metabolites **(Figure 1F, S1H-O**, **2A, B)**. Between spike protein treatments there was some variation, with CoV-2 inducing rhythmic changes to the TCA cycle and its metabolites and OXPHOS proteins **(Figure 4A, S4A, B)**. Spike protein treatment at CT16-20 had the opposite effect on central metabolism. At CT16-20, both glycolysis and TCA cycle pathways are at their nadir and OXPHOS is at its highest relative abundance. Here, spike protein exposure upregulated both glycolysis and the TCA cycle, with little effect on OXPHOS. In total, these data suggest spike protein exposure further suppresses immunometabolism during the anti-inflammatory phase while amplifying it during the pro-inflammatory phase.

Given the widespread temporal changes to immunometabolic pathways, we hypothesized that mitochondrial function and morphology would be similarly altered by spike protein exposure. Mitochondria serve as key regulators of the TCA cycle and OXPHOS pathways, and aspects of their morphology and function are coordinated by circadian rhythms [58]. We found mitochondrial membrane potential (MMP)—a readout of OXPHOS function—oscillated with a circadian period and peaked between CT10-14, just prior to the peak of OXPHOS proteins at CT18 **(Figure S5A, Figure 1F)**. Treatment with spike protein at CT8 decreased MMP levels, correlating with the downregulation of OXPHOS proteins and cardiolipins **(Figure 4B)**. Cardiolipins are exclusive to mitochondria and anchor OXPHOS complexes to the inner membrane, and their loss leads to downregulation of OXPHOS proteins and an inability to drive pro-inflammatory responses in mBMDMs [87]. This suggests the downregulation we observed at CT8 contributed to decreased OXPHOS proteins and MMP at that time. Mitochondria also exhibited increased branching at this time, suggesting dynamic changes to their morphology and deviation from their innate fission-fusion rhythms [10]. Although the role of increased mitochondrial branching is unclear, *in silico* models suggest it could be indicative of attempts to increase network formation to recover normal mitochondrial metabolic function [88]. Intriguingly, the spike protein induced changes to MMP and morphology do not mimic those of the classical stimuli (e.g. LPS), further indicating that spike protein induces different responses than seen in classical models of inflammation.

In parallel with what we noted in mice, human macrophages exhibited a similar response to spike proteins. We observed little classical inflammatory response but instead found rhythmic responses to spike proteins for proteins within central metabolic pathways and mitochondria. However, we did find nuanced differences between the two species. In humans, the immunometabolic repression was evident around HPS20-24. Additionally, in humans, several components of the immune response, including BMAL1, macropinocytosis proteins, and the ROS scavenger SOD2, were all downregulated at this time, suggesting an overall decrease in ability to properly internalize and process spike protein at HPS20-24 **(Figure 6B, Supplemental Figure S8G, H)** [89,90]. An increased metabolic response in humans occurred closer to HPS32-36 and was characterized by an increase in glycolysis but a corresponding decrease in TCA proteins. Together, these data suggest that spike protein exposure in both species evokes a modest inflammatory response, timed by the circadian control over immunometabolism.

There are several potential limitations to this work. The mice used in these experiments did not contain the humanized ACE2 receptor gene and thus may have low binding efficiency between ACE2 and WT spike protein [91]. However, current literature suggests that stimulation by toll-like receptors TLR2 and TLR4 may drive macrophage responses, and increasing mutations in spike protein variants appear to bind more tightly to mouse ACE2 [91–94]. Additionally, given the similarities in immunometabolic responses we observed between mouse and humans, we propose that our findings reflect conserved pathways that are relevant to spike protein exposure. Further, our analysis of spike protein exposure was 6 hours long, to capture circadian effects. Longer exposure and *in vivo* studies would be necessary to ascertain whether the initial trends we described persist in long term effects.

Overall, our analysis demonstrates that exposure to CoV spike proteins 1) does have a time-of-day effect and 2) that this effect does not tend to involve the classical markers of an inflammatory response. Instead, most temporal differences in the response to CoV spike proteins were within central metabolism and mitochondrial function, which aligns with the circadian regulation of immunometabolism. While there were some notable differences between humans and mice, in the aggregate, the response was similar, suggesting that the response of macrophages to the spike protein is conserved across species. Moreover, although the intensity of ACE2 binding may differ between the two, the tightness of the binding may not be the only factor in eliciting a response. We predict that, given the similarities in CoV responses across cell types in the literature and widespread circadian regulation of cellular metabolism, the circadian immunometabolic response likely extends to other cell types [9,16,17,43,95].

The temporal responses we identified in this work may help to explain the underlying biology behind the rhythms in COVID vaccine efficacy based on the time of day of administration. In human trials, COVID vaccination in the morning was positively associated with less risk of severe COVID infection, a pattern that was found with several other vaccines [14,96]. This suggests the involvement of daily immune rhythms in vaccination response. Given their role as the immune system’s first responders, macrophages are a key cell type in early vaccination response [97]. With other studies demonstrating rhythms in the adaptive immune system that parallel what is seen in macrophages, our data suggest that the circadian timing of immunometabolism is the driving factor in this response **(Figure 7)** [98].

## Acknowledgements

We thank Dr. Antigone McKenna and the staff in the Rensselaer BioResearch Core for animal care and Dr. Sergey Pryschep of the Rensselaer Microscopy Core for imaging expertise. We also thank Cameron R. Plowinske for computational support and all members of the Hurley Lab at RPI for their valuable feedback and support. Finally, we gratefully acknowledge the anonymous human donor whose contribution made this research possible.

## Supporting Information

**S1 Fig.** Macrophage proteomic, metabolomic, and lipidomic data demonstrate highly correlated data and circadian oscillations in core clock proteins and central metabolic pathways. Control = blue CoV-1 = teal, CoV-2 = green, CoV-2^D614G^ = yellow. **(A)** A PCA analysis of the proteomics data, depicting variation of each time point and replicate on PC1 and PC2. **(B)** Correlation plots of each proteomic replicate compared to all proteomics replicates at each time point, separated by treatment type (indicated by the box color) with highly positive correlations shown in orange. **(C)** A PCA analysis of the metabolic data, depicting variation of each time point and replicate on PC1 and PC2. **(D)** Correlation plots of each metabolic replicate compared to all metabolic replicates at each time point, separated by treatment type (indicated by the box color) with highly positive correlations shown in orange. **(E)** A PCA analysis of the lipidomic data, depicting variation of each time point and replicate on PC1 and PC2. **(F)** Correlation plots of each lipidomic replicate compared to all lipidomics replicates at each time point, separated by treatment type (indicated by the box color) with highly positive correlations shown in orange. **(G)** Lines reflect ECHO fitted models for core circadian clock proteins detected in the proteomics data set. Shading depicts ±1 standard deviation of the data. **(H-K)** Abundance values (points) and ECHO-fitted models (line) with statistical significance for all detected central metabolism proteins in **(H)** glycolysis, **(I)** pentose phosphate pathway, **(J)** TCA cycle, and **(K)** OXPHOS. **(L-M)** ECHO fit of circadian OXPHOS proteins in Complex I **(L)**, Complex III **(M)**, Complex IV **(N)**, and Complex V **(O)**. Statistical significance was determined using the Friedman test and post-hoc Dunn’s Test, * = p<0.05, ** = p<0.01, ***= p<0.001, ****= p<0.0001

**S2 Fig.** Spike protein exposure does not affect oscillations of PER2. **(A)** The percent similarity of peptide chains between all spike proteins as determined by MOE (2024.06). **(B)** Lumicycle traces tracking PER2::LUC luminescence counts after exposure to spike proteins at HPS16, 20, 24, 28, 32, 36 (dotted red line denotes exposure time). **(C)** Circadian period difference from control at each time point after exposure to spike proteins. Error bars represent ±1SD from the mean. **(D)** Lumicycle traces tracking PER2::LUC luminescence counts after exposure to LPS and IL-4 at HPS16, 20, 24, 28, 32, 36 (dotted red line denotes exposure time). **(E)** Differential levels of all detected circadian clock proteins after exposure to spike proteins. Gray lines indicate log2FC and FDR significance cutoffs. Clock proteins are represented by shapes and spike protein treatments are indicated by colors. Statistics: RM 2-way ANOVA with Geisser-Greenhouse correction and post-hoc Tukey’s multiple comparison test. ns = not significant.

**S3 Fig.** Macrophage response to spike proteins affects central metabolism in a circadian manner. Cytokine levels in response to vehicle control, CoV-1, CoV-2, or CoV-2^D614G^ spike protein for **(A)** IL-6, **(B)** TNFa, **(C)** TGFβ-1, and **(D)** IL-10. **(E)** TGF-*β*1 cytokine levels normalized to vehicle control levels at each time point of exposure to CoV-1 spike protein (ratio of CoV-1:vehicle control represented as bars), with significant ECHO fitted model (line). Cytokine levels in response to treatment with LPS and IL-4 for **(F)** TGF-*β*1 and **(G)** IL-10. HPS = hours post serum shock, CT = circadian time. **(H-K)** Violin plots showing all protein log2FC values (each point represents one protein log2FC value) and significant circadian ECHO models (lines) for response to spike proteins in **(H)** glycolysis, **(I)** pentose phosphate pathway (PPP), **(J)** TCA, and **(K)** OXPHOS pathways. Statistical significance for cytokines determined using RM 2-way ANOVA with Geisser-Greenhouse correction and post-hoc Dunnett’s multiple comparison test. Statistical significance for central metabolic pathway violin plots determined using Kruskall-Wallis and post-hoc Dunn’s Test. * = p<0.05, ** = p<0.01, *** = p<0.001, **** = p<0.000.

**S4 Fig.** Mitochondrial terms and proteins are altered upon spike protein exposure. Significant GO ontologies (PantherGO) of mitochondrial-related significant DEPs in response to CoV-2^D614G^ spike protein. **(B)** A chart of all significant mitochondrial DEPs (MitoCarta 3.0[44]) in response to treatment by CoV-1, CoV-2, and CoV-2^D614G^ spike proteins (colors in key) at HPS16, 20, 24, 28, 32, 36, and 40. Internal numbers represent the fold increase (light) or decrease (dark) relative to PBS control.

**S5 Fig.** Mitochondrial membrane potential oscillates with a 24-hr period and is altered upon treatment with pro- and anti-inflammatory stimuli. **(A)** Measurements of JC-10 590/525 ratios under non-stimulatory PBS control conditions across 24 hrs. Line represents ECHO model filtered for circadian parameters (period = 20-28hrs, BH-adj p-value = <0.05). **(B)** Mitochondrial membrane potential (MMP) at each treatment time point, as a percentage of the vehicle control, for CoV-1, CoV-2, and CoV-2^D614G^ spike protein treatments. **(C)** JC-10 590/525 ratios of mBMDMs treated at HPS16, 20, 24, 28, 32, 36, or 40 with LPS (left) or IL-4 (right), as a percentage of the PBS-treated control. Overlaid line on all graphs indicates significant ECHO fit if response was determined to be circadian (e.g. period = 20-28h, BH-adj p-val = <0.05). **(A-C)** Bar colors indicate treatment type, and individual point colors represent each biological replicate as indicated in the key. Error bars represent ±1 SD. **(D)** Confocal maximum projections of representative fixed mBMDMs stained with MitoTracker^TM^ CMXRos Red following treatment with either PBS control, CoV-2 spike protein, LPS, or IL-4 at HPS20, 24, 32, and 36. Images pseudo-colored using the inverted look-up table (LUT) in ImageJ/FIJI. Omitted images indicated within the panels are reported in Figure 4C. Scale bar represents 2μM.

**S6 Fig.** Human proteomics, metabolomics, and lipidomics samples vary by treatment and cluster by time of day. **(A)** PCA analysis showing variation between control and CoV-2 spike-treated samples on PC1 and PC2 axes, colored by treatment type. **(B)** PCA analysis showing variation of only PBS control samples, colored by time of day, and plotted on PC1 and PC2 axes. **(C)** PCA analysis of the metabolomic data, depicting variation of each time point and replicate on PC1 and PC2. **(D)** PCA analyses of the lipidomic data, depicting variation of each time point and replicate on PC1 and PC2 axes, colored by treatment type and separated by positive and negative MS acquisition modes. **(E)** Correlation plots of each metabolomic replicate compared to all metabolomic replicates at each time point, separated by treatment type (indicated by the box color). **(F)** Correlation plots of each lipidomic replicate compared to all lipidomic replicates at each time point, separated by MS positive and negative acquisition mode and treatment type (indicated by the box color). Control = dark blue, CoV-2 = light blue, CoV-2α = salmon.

**S7 Fig.** The circadian clock coordinates peak timing of multiple lipid classes, mitochondrial proteins, and key central metabolic processes in human macrophages. **(A)** Heatmaps representing ECHO models of circadian proteins within the context of the glycolysis pathway. **(B)** Heatmaps representing ECHO models of circadian proteins within the context of the TCA cycle. **(C)** Heatmaps representing ECHO models of circadian proteins within the complexes of oxidative phosphorylation (OXPHOS). **(D)** Percent of circadian lipids within each main lipid class. Number of circadian/total in class for each lipid class is listed after each bar. Acylcarnitines (CAR), dihexosylceramides (Hex2Cer), diacylglycerols (DG), phosphatidylglycerols (PG), cholesteryl esters (CE), phosphatidylethanolamines (PE), ceramides/hexosylceramides (CE/HexCer), phosphatidylcholines (PC), phosphatidylserines (PS), sphingomyelins (SM), triglycerides (TG), phosphatidylinositols (PI), **(E)** ECHO fit models (line) of circadian Phosphatidylinositols (PI). Shaded portion represents ±1SD of the data. **(F)** Peak timing of circadian mitochondrial proteins, grouped by MitoCarta3.0 annotation [44]. Important sub-processes and proteins indicated by text and brackets.

**S8 Fig.** Metabolic pathways in human macrophages respond in a time-of day dependent manner to CoV-2α. **(A-B)** Bar graphs plotting the number of up- and down-regulated proteins at each HPS time point following treatment with either **(A)** CoV-2 or CoV-2α spike protein. **(C)** Volcano plot of differential levels of detected core clock proteins when treated at each time point with CoV-2α spike protein. Symbol shape indicates clock protein; color indicates treatment time point (HPS); dotted lines indicate significant log2FC and -log10(FDR) cutoff values. **(D)** ECHO model fit lines to log2FC values of all central metabolic proteins, separated by pathway, in response to CoV-2α spike protein. Significant fits are denoted by (*). **(E)** Bubble plot of all detected metabolites within the central metabolic pathways glycolysis and TCA cycle in response to CoV-2α spike protein. Color indicates significant differential levels, bubble size denotes –log10 FDR value. Metabolites are listed in the order they follow in each pathway. **(F)** Violin plots showing all protein log2FC values (each point represents one protein log2FC value) and significant circadian ECHO models (lines) for glycolysis, TCA cycle, and OXPHOS in response to CoV-2 (blue) and CoV-2α (salmon pink). **(G)** Line graph showing ECHO fit to log2FC of mitochondrial-related circadian response proteins over circadian time (HPS), in response to CoV-2 spike protein. **(H)** Line graph showing ECHO fit to log2FC of plasma membrane organization-related circadian response proteins over circadian time (HPS) in response to CoV-2 spike protein.

## References

1. Sender R, Weiss Y, Navon Y, Milo I, Azulay N, Keren L, et al. The total mass, number, and distribution of immune cells in the human body. Proc Natl Acad Sci. 2023;120: e2308511120. doi:10.1073/pnas.2308511120

2. Langston PK, Shibata M, Horng T. Metabolism Supports Macrophage Activation. Front Immunol. 2017;8. doi:10.3389/fimmu.2017.00061

3. Li J, Shan R, Miller H, Filatov A, Byazrova MG, Yang L, et al. The roles of macrophages and monocytes in COVID-19 Severe Respiratory Syndrome. Cell Insight. 2025;4: 100250. doi:10.1016/j.cellin.2025.100250

4. Simonis A, Theobald SJ, Koch AE, Mummadavarapu R, Mudler JM, Pouikli A, et al. Persistent epigenetic memory of SARS-CoV-2 mRNA vaccination in monocyte-derived macrophages. Mol Syst Biol. 2025;21: 341–360. doi:10.1038/s44320-025-00093-6

5. Mamilos A, Winter L, Schmitt VH, Barsch F, Grevenstein D, Wagner W, et al. Macrophages: From Simple Phagocyte to an Integrative Regulatory Cell for Inflammation and Tissue Regeneration—A Review of the Literature. Cells. 2023;12: 276. doi:10.3390/cells12020276

6. Wculek SK, Dunphy G, Heras-Murillo I, Mastrangelo A, Sancho D. Metabolism of tissue macrophages in homeostasis and pathology. Cell Mol Immunol. 2022;19: 384–408. doi:10.1038/s41423-021-00791-9

7. Mills EL, Kelly B, Logan A, Costa ASH, Varma M, Bryant CE, et al. Repurposing mitochondria from ATP production to ROS generation drives a pro-inflammatory phenotype in macrophages that depends on succinate oxidation by complex II. Cell. 2016;167: 457–470.e13. doi:10.1016/j.cell.2016.08.064

8. Mills EL, Kelly B, O’Neill LAJ. Mitochondria are the powerhouses of immunity. Nat Immunol. 2017;18: 488–498. doi:10.1038/ni.3704

9. Lin Y, He L, Cai Y, Wang X, Wang S, Li F. The role of circadian clock in regulating cell functions: implications for diseases. MedComm. 2024;5: e504. doi:10.1002/mco2.504

10. Collins EJ, Cervantes-Silva MP, Timmons GA, O’Siorain JR, Curtis AM, Hurley JM. Post-transcriptional circadian regulation in macrophages organizes temporally distinct immunometabolic states. Genome Res. 2021;31: 171–185. doi:10.1101/gr.263814.120

11. Keller M, Mazuch J, Abraham U, Eom GD, Herzog ED, Volk H-D, et al. A circadian clock in macrophages controls inflammatory immune responses. Proc Natl Acad Sci U S A. 2009;106: 21407–21412. doi:10.1073/pnas.0906361106

12. Ramond E, Jamet A, Coureuil M, Charbit A. Pivotal Role of Mitochondria in Macrophage Response to Bacterial Pathogens. Front Immunol. 2019;10: 2461. doi:10.3389/fimmu.2019.02461

13. Rurek M. Mitochondria in COVID-19: from cellular and molecular perspective. Front Physiol. 2024;15. doi:10.3389/fphys.2024.1406635

14. Hazan G, Duek OA, Alapi H, Mok H, Ganninger A, Ostendorf E, et al. Biological rhythms in COVID-19 vaccine effectiveness in an observational cohort study of 1.5 million patients. J Clin Invest. 2023;133. doi:10.1172/JCI167339

15. Jackson CB, Farzan M, Chen B, Choe H. Mechanisms of SARS-CoV-2 entry into cells. Nat Rev Mol Cell Biol. 2022;23: 3–20. doi:10.1038/s41580-021-00418-x

16. Clough E, Inigo J, Chandra D, Chaves L, Reynolds JL, Aalinkeel R, et al. Mitochondrial Dynamics in SARS-COV2 Spike Protein Treated Human Microglia: Implications for Neuro-COVID. J Neuroimmune Pharmacol. 2021;16: 770–784. doi:10.1007/s11481-021-10015-6

17. Cao X, Nguyen V, Tsai J, Gao C, Tian Y, Zhang Y, et al. The SARS-CoV-2 spike protein induces long-term transcriptional perturbations of mitochondrial metabolic genes, causes cardiac fibrosis, and reduces myocardial contractile in obese mice. Mol Metab. 2023;74: 101756. doi:10.1016/j.molmet.2023.101756

18. Huynh TV, Rethi L, Lee T-W, Higa S, Kao Y-H, Chen Y-J. Spike Protein Impairs Mitochondrial Function in Human Cardiomyocytes: Mechanisms Underlying Cardiac Injury in COVID-19. Cells. 2023;12: 877. doi:10.3390/cells12060877

19. Duan L, Zheng Q, Zhang H, Niu Y, Lou Y, Wang H. The SARS-CoV-2 Spike Glycoprotein Biosynthesis, Structure, Function, and Antigenicity: Implications for the Design of Spike-Based Vaccine Immunogens. Front Immunol. 2020;11. doi:10.3389/fimmu.2020.576622

20. Buel SM, Debopadhaya S, De Los Santos H, Edwards KM, David AM, Dao UH, et al. The PAICE suite reveals circadian posttranscriptional timing of noncoding RNAs and spliceosome components in Mus musculus macrophages. G3. 2022;12: jkac176. doi:10.1093/g3journal/jkac176

21. Hughes ME, Abruzzi KC, Allada R, Anafi R, Arpat AB, Asher G, et al. Guidelines for Genome-Scale Analysis of Biological Rhythms. J Biol Rhythms. 2017;32: 380–393. doi:10.1177/0748730417728663

22. Nakayasu ES, Nicora CD, Sims AC, Burnum-Johnson KE, Kim Y-M, Kyle JE, et al. MPLEx: a Robust and Universal Protocol for Single-Sample Integrative Proteomic, Metabolomic, and Lipidomic Analyses. mSystems. 2016;1: e00043–16. doi:10.1128/mSystems.00043-16

23. De los Santos H, Collins EJ, Mann C, Sagan AW, Jankowski MS, Bennett KP, et al. ECHO: an application for detection and analysis of oscillators identifies metabolic regulation on genome-wide circadian output. Kelso J, editor. Bioinformatics. 2020;36: 773–781. doi:10.1093/bioinformatics/btz617

24. Crowell AM, Greene CS, Loros JJ, Dunlap JC. Learning and Imputation for Mass-spec Bias Reduction (LIMBR). Bioinformatics. 2019;35: 1518–1526. doi:10.1093/bioinformatics/bty828

25. Kanehisa M, Furumichi M, Sato Y, Matsuura Y, Ishiguro-Watanabe M. KEGG: biological systems database as a model of the real world. Nucleic Acids Res. 2024;53: D672–D677. doi:10.1093/nar/gkae909

26. Zhang H, Jay Forman H, Choi J. γ-Glutamyl Transpeptidase in Glutathione Biosynthesis. In: Sies H, Packer L, editors. Methods in Enzymology. Academic Press; 2005. pp. 468–483. doi:10.1016/S0076-6879(05)01028-1

27. KEGG COMPOUND: C01879. [cited 19 May 2025]. Available: https://www.genome.jp/dbget-bin/www_bget?C01879

28. Shakespear MR, Iyer A, Cheng CY, Das Gupta K, Singhal A, Fairlie DP, et al. Lysine Deacetylases and Regulated Glycolysis in Macrophages. Trends Immunol. 2018;39: 473–488. doi:10.1016/j.it.2018.02.009

29. Lu M, Luo D, Zhang Z, Ouyang F, Shi Y, Hu C, et al. Branched-chain amino acid catabolism promotes M2 macrophage polarization. Front Immunol. 2024;15: 1469163. doi:10.3389/fimmu.2024.1469163

30. Tang Y, Yu Y, Li R, Tao Z, Zhang L, Wang X, et al. Phenylalanine promotes alveolar macrophage pyroptosis via the activation of CaSR in ARDS. Front Immunol. 2023;14. doi:10.3389/fimmu.2023.1114129

31. Decker ST, Funai K. Mitochondrial membrane lipids in the regulation of bioenergetic flux. Cell Metab. 2024;36: 1963–1978. doi:10.1016/j.cmet.2024.07.024

32. Rutkowsky JM, Knotts TA, Ono-Moore KD, McCoin CS, Huang S, Schneider D, et al. Acylcarnitines activate proinflammatory signaling pathways. Am J Physiol Endocrinol Metab. 2014;306: E1378–1387. doi:10.1152/ajpendo.00656.2013

33. Paradies G, Paradies V, Ruggiero FM, Petrosillo G. Role of Cardiolipin in Mitochondrial Function and Dynamics in Health and Disease: Molecular and Pharmacological Aspects. Cells. 2019;8: 728. doi:10.3390/cells8070728

34. Sievers BL, Cheng MTK, Csiba K, Meng B, Gupta RK. SARS-CoV-2 and innate immunity: the good, the bad, and the “goldilocks.” Cell Mol Immunol. 2024;21: 171–183. doi:10.1038/s41423-023-01104-y

35. Robinson MD, McCarthy DJ, Smyth GK. edgeR: a Bioconductor package for differential expression analysis of digital gene expression data. Bioinformatics. 2010;26: 139–140. doi:10.1093/bioinformatics/btp616

36. Szklarczyk D, Kirsch R, Koutrouli M, Nastou K, Mehryary F, Hachilif R, et al. The STRING database in 2023: protein–protein association networks and functional enrichment analyses for any sequenced genome of interest. Nucleic Acids Res. 2023;51: D638–D646. doi:10.1093/nar/gkac1000

37. Chen S, Fuller KK, Dunlap JC, Loros JJ. A Pro- and Anti-inflammatory Axis Modulates the Macrophage Circadian Clock. Front Immunol. 2020;11: 867. doi:10.3389/fimmu.2020.00867

38. Lellupitiyage Don SS, Mas-Rosario JA, Lin H-H, Nguyen EM, Taylor SR, Farkas ME. Macrophage circadian rhythms are differentially affected based on stimuli. Integr Biol. 2022;14: 62–75. doi:10.1093/intbio/zyac007

39. Francisco JC, Virshup DM. Casein Kinase 1 and Human Disease: Insights From the Circadian Phosphoswitch. Front Mol Biosci. 2022;9: 911764. doi:10.3389/fmolb.2022.911764

40. Zhao B, Nepovimova E, Wu Q. The role of circadian rhythm regulator PERs in oxidative stress, immunity, and cancer development. Cell Commun Signal CCS. 2025;23: 30. doi:10.1186/s12964-025-02040-2

41. Rodrigues TS, Zamboni DS. Inflammasome activation by SARS-CoV-2 and its participation in COVID-19 exacerbation. Curr Opin Immunol. 2023;84: 102387. doi:10.1016/j.coi.2023.102387

42. O’Neill LAJ, Kishton RJ, Rathmell J. A guide to immunometabolism for immunologists. Nat Rev Immunol. 2016;16: 553–565. doi:10.1038/nri.2016.70

43. Kulkovienė G, Narauskaitė D, Tunaitytė A, Volkevičiūtė A, Balion Z, Kutakh O, et al. Differential Mitochondrial, Oxidative Stress and Inflammatory Responses to SARS-CoV-2 Spike Protein Receptor Binding Domain in Human Lung Microvascular, Coronary Artery Endothelial and Bronchial Epithelial Cells. Int J Mol Sci. 2024;25: 3188. doi:10.3390/ijms25063188

44. Rath S, Sharma R, Gupta R, Ast T, Chan C, Durham TJ, et al. MitoCarta3.0: an updated mitochondrial proteome now with sub-organelle localization and pathway annotations. Nucleic Acids Res. 2021;49: D1541–D1547. doi:10.1093/nar/gkaa1011

45. Angireddy R, Kazmi HR, Srinivasan S, Sun L, Iqbal J, Fuchs SY, et al. Cytochrome c oxidase dysfunction enhances phagocytic function and osteoclast formation in macrophages. FASEB J Off Publ Fed Am Soc Exp Biol. 2019;33: 9167–9181. doi:10.1096/fj.201900010RR

46. Kunji ERS, Aleksandrova A, King MS, Majd H, Ashton VL, Cerson E, et al. The transport mechanism of the mitochondrial ADP/ATP carrier. Biochim Biophys Acta BBA - Mol Cell Res. 2016;1863: 2379–2393. doi:10.1016/j.bbamcr.2016.03.015

47. Masquelier J, Alhouayek M, Terrasi R, Bottemanne P, Paquot A, Muccioli GG. Lysophosphatidylinositols in inflammation and macrophage activation: Altered levels and anti-inflammatory effects. Biochim Biophys Acta BBA - Mol Cell Biol Lipids. 2018;1863: 1458–1468. doi:10.1016/j.bbalip.2018.09.003

48. Oliva-Ramírez J, Moreno-Altamirano MMB, Pineda-Olvera B, Cauich-Sánchez P, Sánchez-García FJ. Crosstalk between circadian rhythmicity, mitochondrial dynamics and macrophage bactericidal activity. Immunology. 2014;143: 490–497. doi:10.1111/imm.12329

49. Mishra P, Chan DC. Metabolic regulation of mitochondrial dynamics. J Cell Biol. 2016;212: 379–387. doi:10.1083/jcb.201511036

50. Shin HJ, Lee W, Ku KB, Yoon GY, Moon H-W, Kim C, et al. SARS-CoV-2 aberrantly elevates mitochondrial bioenergetics to induce robust virus propagation. Signal Transduct Target Ther. 2024;9: 125. doi:10.1038/s41392-024-01836-x

51. Afroz SF, Raven KD, Lawrence GMEP, Kapetanovic R, Schroder K, Sweet MJ. Mitochondrial dynamics in macrophages: divide to conquer or unite to survive? Biochem Soc Trans. 2023;51: 41–56. doi:10.1042/BST20220014

52. Chaudhry A, Shi R, Luciani DS. A pipeline for multidimensional confocal analysis of mitochondrial morphology, function, and dynamics in pancreatic β-cells. Am J Physiol - Endocrinol Metab. 2020;318: E87–E101. doi:10.1152/ajpendo.00457.2019

53. Dill-McFarland KA, Peterson G, Lim PN, Skerrett S, Hawn TR, Rothchild AC, et al. Shared and distinct responses of human and murine alveolar macrophages and monocyte-derived macrophages to Mycobacterium tuberculosis. ImmunoHorizons. 2025;9: vlaf051. doi:10.1093/immhor/vlaf051

54. Tanaka K, Takenaka S, Yoshida K. Scyllo-Inositol, a Therapeutic Agent for Alzheimer’s Disease. Austin J Clin Neurol. 2015;2. Available: https://austinpublishinggroup.com/clinical-neurology/fulltext/ajcn-v2-id1040.pdf

55. López-Gambero AJ, Sanjuan C, Serrano-Castro PJ, Suárez J, Rodríguez de Fonseca F. The Biomedical Uses of Inositols: A Nutraceutical Approach to Metabolic Dysfunction in Aging and Neurodegenerative Diseases. Biomedicines. 2020;8: 295. doi:10.3390/biomedicines8090295

56. O’Donnell VB, Rossjohn J, Wakelam MJO. Phospholipid signaling in innate immune cells. J Clin Invest. 128: 2670–2679. doi:10.1172/JCI97944

57. Balla T. Phosphoinositides: Tiny Lipids With Giant Impact on Cell Regulation. Physiol Rev. 2013;93: 1019–1137. doi:10.1152/physrev.00028.2012

58. Marques E, Kramer R, Ryan DG. Multifaceted mitochondria in innate immunity. Npj Metab Health Dis. 2024;2: 6. doi:10.1038/s44324-024-00008-3

59. Merrill RA, Strack S. MITOCHONDRIA: A kinase anchoring protein 1, a signaling platform for mitochondrial form and function. Int J Biochem Cell Biol. 2014;48: 92–96. doi:10.1016/j.biocel.2013.12.012

60. Wang N, Wang X, Lan B, Gao Y, Cai Y. DRP1, fission and apoptosis. Cell Death Discov. 2025;11: 150. doi:10.1038/s41420-025-02458-0

61. Lazarevic I, Pravica V, Miljanovic D, Cupic M. Immune Evasion of SARS-CoV-2 Emerging Variants: What Have We Learnt So Far? Viruses. 2021;13: 1192. doi:10.3390/v13071192

62. Richter-Dennerlein R, Korwitz A, Haag M, Tatsuta T, Dargazanli S, Baker M, et al. DNAJC19, a Mitochondrial Cochaperone Associated with Cardiomyopathy, Forms a Complex with Prohibitins to Regulate Cardiolipin Remodeling. Cell Metab. 2014;20: 158–171. doi:10.1016/j.cmet.2014.04.016

63. Janz A, Walz K, Cirnu A, Surjanto J, Urlaub D, Leskien M, et al. Mutations in DNAJC19 cause altered mitochondrial structure and increased mitochondrial respiration in human iPSC-derived cardiomyocytes. Mol Metab. 2024;79: 101859. doi:10.1016/j.molmet.2023.101859

64. Barhoumi T, Alghanem B, Shaibah H, Mansour FA, Alamri HS, Akiel MA, et al. SARS-CoV-2 Coronavirus Spike Protein-Induced Apoptosis, Inflammatory, and Oxidative Stress Responses in THP-1-Like-Macrophages: Potential Role of Angiotensin-Converting Enzyme Inhibitor (Perindopril). Front Immunol. 2021;12. doi:10.3389/fimmu.2021.728896

65. Palma FR, He C, Danes JM, Paviani V, Coelho DR, Gantner BN, et al. Mitochondrial Superoxide Dismutase: What the Established, the Intriguing, and the Novel Reveal About a Key Cellular Redox Switch. Antioxid Redox Signal. 2020;32: 701–714. doi:10.1089/ars.2019.7962

66. Ahn W, Burnett FN, Pandey A, Ghoshal P, Singla B, Simon AB, et al. SARS-CoV-2 Spike Protein Stimulates Macropinocytosis in Murine and Human Macrophages via PKC-NADPH Oxidase Signaling. Antioxidants. 2024;13: 175. doi:10.3390/antiox13020175

67. Zhang Y-Y, Liang R, Wang S-J, Ye Z-W, Wang T-Y, Chen M, et al. SARS-CoV-2 hijacks macropinocytosis to facilitate its entry and promote viral spike–mediated cell-to-cell fusion. J Biol Chem. 2022;298: 102511. doi:10.1016/j.jbc.2022.102511

68. Yoo S-H, Yamazaki S, Lowrey PL, Shimomura K, Ko CH, Buhr ED, et al. PERIOD2::LUCIFERASE real-time reporting of circadian dynamics reveals persistent circadian oscillations in mouse peripheral tissues. Proc Natl Acad Sci. 2004;101: 5339–5346. doi:10.1073/pnas.0308709101

69. Gerber A, Esnault C, Aubert G, Treisman R, Pralong F, Schibler U. Blood-Borne Circadian Signal Stimulates Daily Oscillations in Actin Dynamics and SRF Activity. Cell. 2013;152: 492–503. doi:10.1016/j.cell.2012.12.027

70. Chemical Computing Group ULC, 910-1010 Sherbrooke St. W., Montreal, QC H3A 2R7. Molecular Operating Environment (MOE), 2024.0601. Available: https://www.chemcomp.com/en/index.htm

71. Monroe ME, Shaw JL, Daly DS, Adkins JN, Smith RD. MASIC: a software program for fast quantitation and flexible visualization of chromatographic profiles from detected LC-MS(/MS) features. Comput Biol Chem. 2008;32: 215–217. doi:10.1016/j.compbiolchem.2008.02.006

72. Kim Y-M, Nowack S, Olsen MT, Becraft ED, Wood JM, Thiel V, et al. Diel metabolomics analysis of a hot spring chlorophototrophic microbial mat leads to new hypotheses of community member metabolisms. Front Microbiol. 2015;6: 209. doi:10.3389/fmicb.2015.00209

73. Kind T, Wohlgemuth G, Lee DY, Lu Y, Palazoglu M, Shahbaz S, et al. FiehnLib: mass spectral and retention index libraries for metabolomics based on quadrupole and time-of-flight gas chromatography/mass spectrometry. Anal Chem. 2009;81: 10038–10048. doi:10.1021/ac9019522

74. Hiller K, Hangebrauk J, Jäger C, Spura J, Schreiber K, Schomburg D. MetaboliteDetector: comprehensive analysis tool for targeted and nontargeted GC/MS based metabolome analysis. Anal Chem. 2009;81: 3429–3439. doi:10.1021/ac802689c

75. Kyle JE, Crowell KL, Casey CP, Fujimoto GM, Kim S, Dautel SE, et al. LIQUID: an-open source software for identifying lipids in LC-MS/MS-based lipidomics data. Bioinforma Oxf Engl. 2017;33: 1744–1746. doi:10.1093/bioinformatics/btx046

76. Conroy MJ, Andrews RM, Andrews S, Cockayne L, Dennis EA, Fahy E, et al. LIPID MAPS: update to databases and tools for the lipidomics community. Nucleic Acids Res. 2023;52: D1677–D1682. doi:10.1093/nar/gkad896

77. R Core Team. R: A Language and Environment for Statistical Computing. Vienna, Austria: R Foundation for Statistical Computing; 2021. Available: https://www.R-project.org/

78. Kanehisa M, Sato Y, Kawashima M, Furumichi M, Tanabe M. KEGG as a reference resource for gene and protein annotation. Nucleic Acids Res. 2016;44: D457–D462. doi:10.1093/nar/gkv1070

79. Thomas PD, Campbell MJ, Kejariwal A, Mi H, Karlak B, Daverman R, et al. PANTHER: A Library of Protein Families and Subfamilies Indexed by Function. Genome Res. 2003;13: 2129–2141. doi:10.1101/gr.772403

80. Searle SR, Speed FM, Milliken GA. Population Marginal Means in the Linear Model: An Alternative to Least Squares Means. Am Stat. 1980;34: 216–221. doi:10.1080/00031305.1980.10483031

81. Kuznetsova A, Brockhoff PB, Christensen RHB. lmerTest Package: Tests in Linear Mixed Effects Models. J Stat Softw. 2017;82: 1–26. doi:10.18637/jss.v082.i13

82. Bates D, Mächler M, Bolker B, Walker S. Fitting Linear Mixed-Effects Models Using lme4. J Stat Softw. 2015;67: 1–48. doi:10.18637/jss.v067.i01

83. Zhang R, Podtelezhnikov AA, Hogenesch JB, Anafi RC. Discovering Biology in Periodic Data through Phase Set Enrichment Analysis (PSEA). J Biol Rhythms. 2016;31: 244–257. doi:10.1177/0748730416631895

84. Gene set enrichment analysis: A knowledge-based approach for interpreting genome-wide expression profiles | PNAS. [cited 24 Oct 2025]. Available: https://www.pnas.org/doi/10.1073/pnas.0506580102

85. Liberzon A, Subramanian A, Pinchback R, Thorvaldsdóttir H, Tamayo P, Mesirov JP. Molecular signatures database (MSigDB) 3.0. Bioinformatics. 2011;27: 1739–1740. doi:10.1093/bioinformatics/btr260

86. Liberzon A, Birger C, Thorvaldsdóttir H, Ghandi M, Mesirov JP, Tamayo P. The Molecular Signatures Database (MSigDB) hallmark gene set collection. Cell Syst. 2015;1: 417–425. doi:10.1016/j.cels.2015.12.004

87. Reynolds MB, Hong HS, Michmerhuizen BC, Lawrence A-LE, Zhang L, Knight JS, et al. Cardiolipin coordinates inflammatory metabolic reprogramming through regulation of Complex II disassembly and degradation. Sci Adv. 9: eade8701. doi:10.1126/sciadv.ade8701

88. Chuphal P, Lanctôt JD, Cornelius SP, Brown AI. Mitochondrial Network Branching Enables Rapid Protein Spread with Slower Mitochondrial Dynamics. PRX Life. 2024;2: 043005. doi:10.1103/PRXLife.2.043005

89. Alexander RK, Liou Y-H, Knudsen NH, Starost KA, Xu C, Hyde AL, et al. Bmal1 integrates mitochondrial metabolism and macrophage activation. Rothlin CV, Rath S, editors. eLife. 2020;9: e54090. doi:10.7554/eLife.54090

90. Palma FR, He C, Danes JM, Paviani V, Coelho DR, Gantner BN, et al. Mitochondrial Superoxide Dismutase: What the Established, the Intriguing, and the Novel Reveal About a Key Cellular Redox Switch. Antioxid Redox Signal. 2020;32: 701–714. doi:10.1089/ars.2019.7962

91. Hatta M, Larson GP, Hatta Y, Wang W, Jiang N, Jung Y-J, et al. ACE2 Receptor Usage across Animal Species by SARS-CoV-2 Variants - Volume 31, Number 8—August 2025 - Emerging Infectious Diseases journal - CDC. 2025 [cited 8 Jan 2026]. doi:10.3201/eid3108.241844

92. Zhao Y, Kuang M, Li J, Zhu L, Jia Z, Guo X, et al. SARS-CoV-2 spike protein interacts with and activates TLR41. Cell Res. 2021;31: 818–820. doi:10.1038/s41422-021-00495-9

93. Khan S, Shafiei MS, Longoria C, Schoggins JW, Savani RC, Zaki H. SARS-CoV-2 spike protein induces inflammation via TLR2-dependent activation of the NF-κB pathway. eLife. 10: e68563. doi:10.7554/eLife.68563

94. Shirato K, Kizaki T. SARS-CoV-2 spike protein S1 subunit induces pro-inflammatory responses via toll-like receptor 4 signaling in murine and human macrophages. Heliyon. 2021;7: e06187. doi:10.1016/j.heliyon.2021.e06187

95. Huynh TV, Rethi L, Lee T-W, Higa S, Kao Y-H, Chen Y-J. Spike Protein Impairs Mitochondrial Function in Human Cardiomyocytes: Mechanisms Underlying Cardiac Injury in COVID-19. Cells. 2023;12: 877. doi:10.3390/cells12060877

96. Otasowie CO, Tanner R, Ray DW, Austyn JM, Coventry BJ. Chronovaccination: Harnessing circadian rhythms to optimize immunisation strategies. Front Immunol. 2022;13: 977525. doi:10.3389/fimmu.2022.977525

97. Damani-Yokota P, Khanna KM. Innate immune memory: The evolving role of macrophages in therapy. eLife. 2025;14: e108276. doi:10.7554/eLife.108276

98. Ince LM, Barnoud C, Lutes LK, Pick R, Wang C, Sinturel F, et al. Influence of circadian clocks on adaptive immunity and vaccination responses. Nat Commun. 2023;14: 476. doi:10.1038/s41467-023-35979-2

